# Biological molybdenum usage stems back to 3.4 billion years ago

**DOI:** 10.1101/2025.04.02.646658

**Authors:** Aya S. Klos, Morgan S. Sobol, Joanne S. Boden, Eva E. Stüeken, Rika E. Anderson, Kurt O. Konhauser, Betül Kaçar

## Abstract

Molybdenum (Mo) is an essential nutrient for most living organisms, serving as a cofactor in a diverse array of molybdoenzymes that catalyze key reactions in several elemental cycles. However, geochemical data suggest that dissolved Mo concentrations in the Archean ocean (before 2.5 billion years ago) were 1-2 orders of magnitude lower than today, raising questions about its bioavailability to early life. Here, we apply a phylogenomic approach to chart the modern biological and environmental distribution of Mo-related enzymes and use phylogenetic reconciliations to reconstruct the evolutionary history of biological Mo usage. Our results reveal the ubiquity of molybdoenzymes across contemporary organisms inhabiting diverse environments. Furthermore, phylogenetic evidence indicates that the earliest molybdoenzymes stem back to the Paleo/Mesoarchean (~3.5-3.0 Gya), facilitating critical energy-harnessing reactions in some of Earth’s most ancient life forms. Taken together, our findings challenge the prevailing view of limited Mo bioavailability on the anoxic early Earth.

## Introduction

The transition metal molybdenum (Mo) displays an enigmatic evolutionary history in biology^1,2^. On one hand, Mo participates in key biogeochemical transformations of carbon (C), nitrogen (N), and sulfur (S)^2^. Its catalytic functions are mediated through incorporation into enzymes via one of two cofactors; the widely utilized molybdopterin cofactor (Moco) and the [Mo:7Fe:9S:C] metallocluster (FeMoco). The latter is uniquely associated with Mo-dependent nitrogenase^2^, the enzyme that catalyzes the reduction of nitrogen gas (N_2_) to ammonia (NH_3_) during a process known as N_2_-fixation, a key step in the global N cycle that constitutes the major source of N to the biosphere. Phylogenetic analyses suggest that nitrogenases date back to the mid-Archean, over 3 billion years ago^3,4^, consistent with geochemical data^5,6^. In contemporary organisms, cellular Mo utilization depends on complex biosynthetic pathways that assemble either Moco or FeMoco prior to their insertion into enzymes^7,8^. These pathways, or alternatives thereof, would have needed to exist in the mid-Archean to enable Mo utilization at that time.

On the other hand, it is believed that the environmental availability of Mo has fluctuated markedly throughout Earth’s history in response to progressive oxygenation of seawater^9,10^. Prior to the origin of oxygenic photosynthesis and the subsequent accumulation of O_2_ in the atmosphere around 2.45 billion years ago^11^ – the Great Oxidation Event (GOE) – dissolved Mo is thought to have been exceedingly low (<5 nM compared to 105 nM in modern seawater^12^). This scarcity would have stemmed from the suppression of oxidative weathering of Mo-bearing sulfide minerals on land, which today represent the primary source of Mo to the oceans^13^. However, a growing body of evidence suggests a high biological demand for Mo in early Earth environments^4–6^, seemingly at odds with the limited Mo supply inferred from geochemical records^12^. Resolving this discrepancy requires new data to better constrain the relationships between Mo availability, microbial metabolism, and environmental redox conditions in the Archean.

In this study, we address this conundrum via an exhaustive examination of Mo uptake, storage, intracellular trafficking, cofactor biosynthesis, and catalysis. In parallel, we examine the history of biological usage of tungsten (W) for transport and catalysis due to shared chemical properties between Mo and W^14^, similarities in the synthesis of their respective cofactors^15^, and studies reporting W replacing Mo in certain enzymes^16^. First, we use phylogenomic tools to map the distribution of Mo/W usage across modern organisms and their ecological context. Second, we reconstruct the evolutionary history of Mo-/W-utilizing proteins across the modern tree of life. Our aim is to identify the earliest biological processes that used Mo, gauge the impact of environmental change on Mo usage, and outline the sequential stages in the evolution of Mo-based enzymes alongside W-based enzymes over time.

## RESULTS

### Mo proteins are widely distributed across life and diverse environments

Molybdoenzymes can be grouped into four major families, excluding nitrogenases. Three of them, namely dimethyl sulfoxide reductases (DMSOR), xanthine oxidases (XO), and sulfite oxidases (SO), incorporate Mo in the form of Moco, which encompasses different structural variants of molybdopterin ligands (Figure 1). A fourth family of enzymes, aldehyde:ferredoxin oxidoreductase (AOR), incorporates a pyranopterin cofactor that instead contains W in place of Mo (termed Wco)^7^. The similarity in chemical properties between Mo and W allows for some degree of interchangeability between these cofactors. Indeed, some DMSOR family enzymes are known to incorporate Wco instead of Moco^17^. The four major enzyme families described above can further be divided into subfamilies based on their specific substrates.

**Figure 1.**
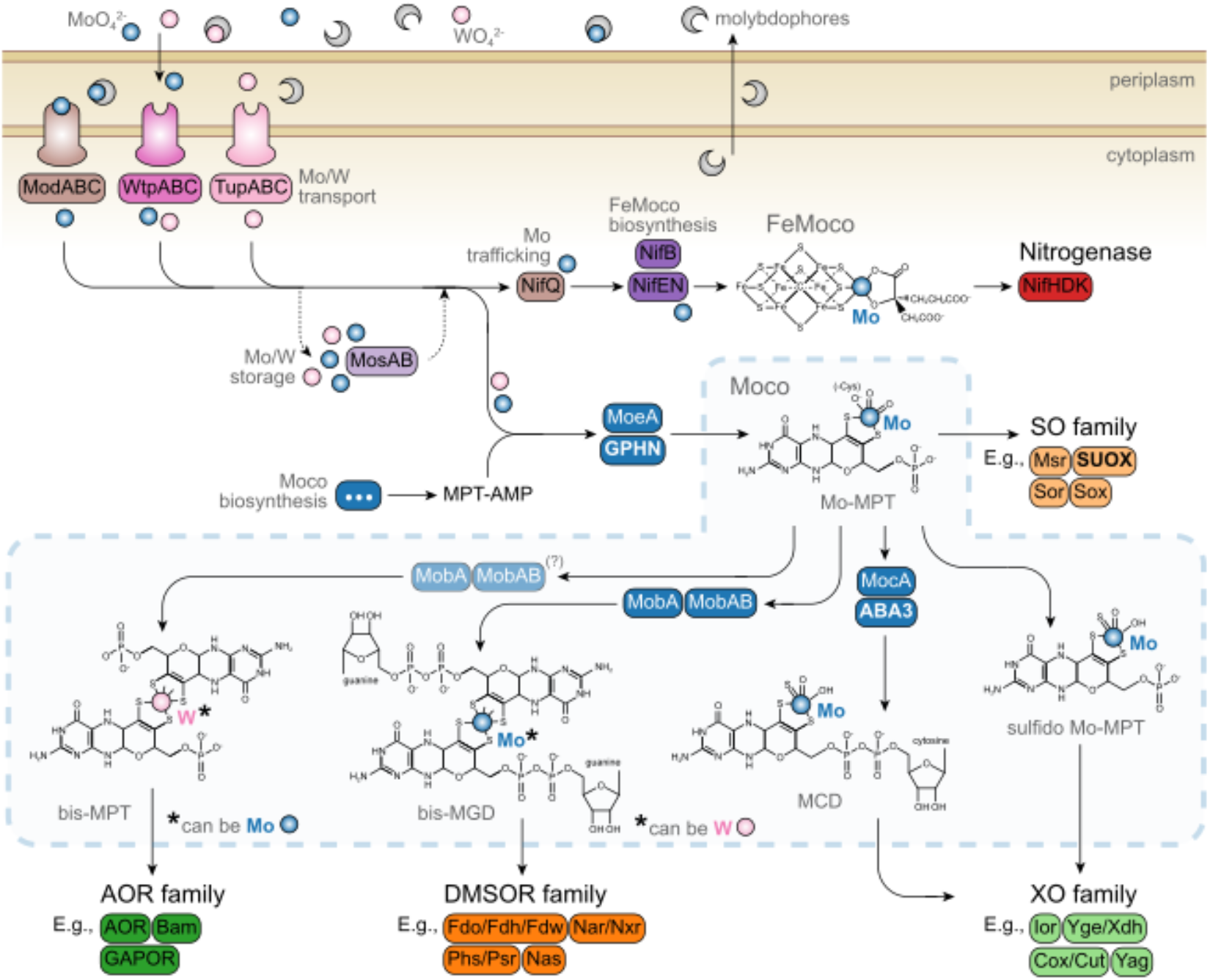
Full schematic of biological Mo-/W-uptake, trafficking and incorporation into selected molybdo-/tungstoenzymes. Chemical structures for the FeMo cofactor (FeMoco) and modified forms of the Mo cofactor (Moco or Mo-MPT for molybdopterin) incorporated by specific enzyme families, as well as a modified form of the W cofactor for AOR family enzymes, are also shown.

In prokaryotes, the incorporation of Mo into molybdoenzymes begins with its uptake from the environment commonly in the form of MoO_4_^2-^ (though biological uptake from Mo-bearing minerals such as molybdenite^18^ has been demonstrated as well). Uptake of this anion by bacteria is most often catalyzed by the high-affinity ABC-type transport system ModABC. In archaea, a dual-affinity ABC-type transport system that is capable of transporting both Mo and W (WtpABC) is reportedly the most common molybdate transporter^19^. Additionally, TupABC, a transporter specifically used for tungstate (WO_4_^-^) uptake, is found in both bacteria and archaea^20^. In eukaryotes, the mechanism for molybdate uptake remains unresolved. However, two different types of molybdate transporters have been identified: MOT1 in land plants and green algae^21,22^, and MOT2 in green algae and animals, including humans^23^. Following uptake, MoO_4_^2-^ undergoes activation and is incorporated into molybdopterin to form Moco. The latter is subsequently incorporated into the enzyme’s active site. For enzymes in the DMSOR, XO, and AOR families, an additional maturation step is required before Moco/Wco incorporation is complete. The core genes involved in Moco biosynthesis pathways are conserved across all domains of life^24^.

We screened select genomes from two previously constructed trees of life^25,26^ to identify genes encompassing the core machinery for Mo(/W) transport, storage, and Mo(/W)co biosynthesis (see methods for protein selection). We also included genes representing all four molybdo/tungstoenzyme families, as well as those involved in nitrogenase and FeMoco assembly, resulting in a total of 102 protein groups. Our results demonstrate that Mo-related genes are widely distributed across nearly all major groups of life, encompassing a broad range of ecological conditions, including temperature, O_2_ tolerance, pH, habitat type, and habitat diversity (Figure 2). These environmental parameters display weak correlations with Mo/W genes (R^2^ < 0.1 in all cases, see Figure S2), but generally speaking, organisms with numerous Mo genes are associated with more temperate, alkaline, and oxic environments, whereas organisms with more W genes are associated with hotter, acidic, and anoxic environments (Figure S2).

**Figure 2.**
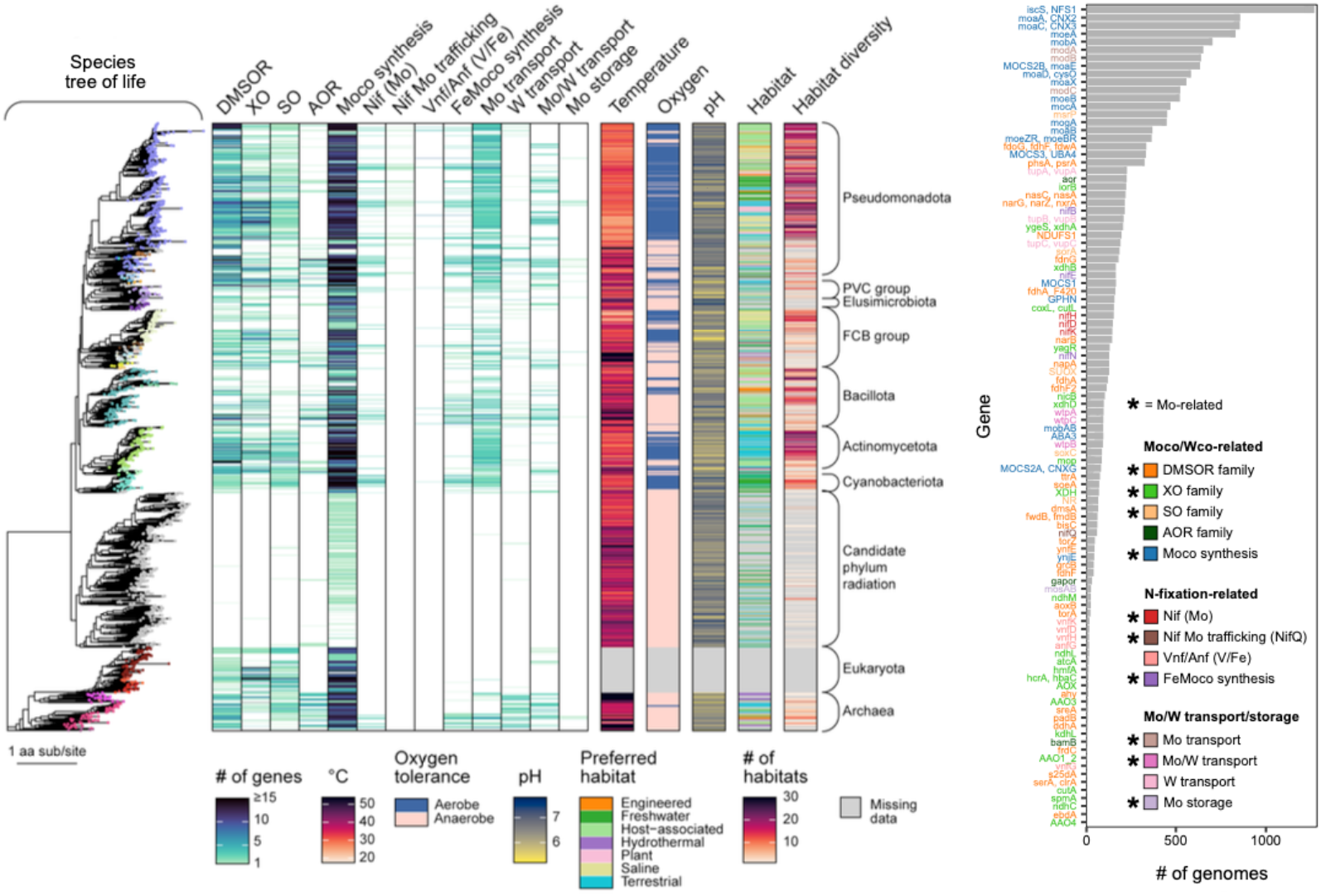
Phylogenetic distribution of Mo-related genes and their associations with host ecological attributes. Mo-related gene counts and host ecological attributes mapped to a species tree of life^25^ (left). Tips on the phylogenetic tree are colored by taxonomy (see Supplementary Figure 1 for color legend. Simplified taxonomic clade labels are on the right. anked frequencies of Mo-related genes across all genomes in our dataset (n = 1,609) (right).

The most widely distributed genes across the analyzed genomes are those involved in Moco biosynthesis, which is essential for the active sites of all molybdoenzymes except nitrogenase. This finding, alongside previous evidence of the pathway’s conservation^24,27^, underscores the fundamental importance of the Moco biosynthesis pathway for Mo- and W-utilizing organisms (W-cofactors rely on the same pathway for synthesis^15^). The next most widely distributed genes encode active membrane transporters for Mo (ModABC) and W (TupABC). The latter is found across nearly all major archaeal lineages but in only select bacterial lineages

(Figure 2). This distribution aligns with previous reports suggesting that Tup transporters are more common in Mo-utilizing archaea compared to bacteria^19^. Among Moco-dependent enzymes, the most widely distributed ones among prokaryotes generally belong to DMSOR. Conversely, the rarest Mo-related genes include certain members of the XO family, W-dependent AOR family, as well as nitrogenase, and Mo-storage genes (MosAB).

### Mo-genes tend to co-occur in extant genomes

To assess the genomic context of Mo-related genes, we analyzed the co-occurrence of individual Mo/W-related protein families across genomes in our dataset (Figure S3). Correlations were also calculated between gene presence and O_2_ preference of genome-associated hosts. Most gene presence correlations are positive, indicating that organisms utilizing Mo typically possess multiple Mo-dependent genes. For example, organisms that use Mo for N_2_-fixation also use Mo for other metabolic pathways and possess the biosynthetic and transport machinery to assemble different types of Mo-cofactors. Given Moco’s flexible redox capacity, which allows it to participate in a wide variety of reactions^1,28^, this finding may reflect a potential evolutionary advantage to maximizing metabolic versatility once the molecular machinery for Mo utilization is established.

W-related genes also co-occur with a few Mo-related genes. This overlap is expected because Wco assembly requires the Moco biosynthetic machinery. However, W-genes also co-occur with high-affinity Mo transporter genes. Notably, many of these co-occurring molybdoenzymes like Fmd and Fdh (used for CO_2_ reduction in the first steps of hydrogenotrophic methanogenesis and formate oxidation, respectively) also have orthologs known to incorporate W^29,30^. These findings suggest an evolutionary advantage to organisms investing in a Mo- and/or W-dependent metabolic system. Once the molecular machinery for Mo/W utilization is established, expanding its use across metabolic pathways may incur little additional cost. This could confer a selective advantage for having dual ability to use Mo and W, particularly in environments where Mo may be limiting, as has been suggested for the early Archean ocean^12^.

### Diversity of Mo-related protein families

We performed a comprehensive assessment of protein similarity among Mo/W-related genes and their homologs (Figure S4). The diverse functions of Mo/W-related proteins are mirrored by their diverse ancestries and origins. We find no detectable sequence homology between most major categories of Mo-related protein families. In other words, they occupy distinct, unconnected clusters within the sequence similarity network. Our analysis also shows no detectable homology between the major Moco/Wco-dependent enzyme families—DMSOR, XO, SO, and AOR—despite them binding chemically similar Mo/W-cofactors. These data suggest that either a shared ancestry is no longer detectable due to the large degree of evolutionary distance or that Moco/Wco-incorporating enzymes evolved independently multiple times throughout life’s history. Even more broadly, the ability of any protein to interact with Mo, regardless of its chemical species, for the purposes of transport, biosynthesis, or catalysis, may have evolved multiple times. Ultimately, this complex history gave rise to distinct enzyme families covering a wide range of substrates and redox potentials. This diversity may reflect the suitability and versatility of Mo as a driving factor for its early incorporation into enzymes on a predominantly anoxic Earth when most Moco and Wco-dependent enzyme families evolved (Figure 3).

**Figure 3.**
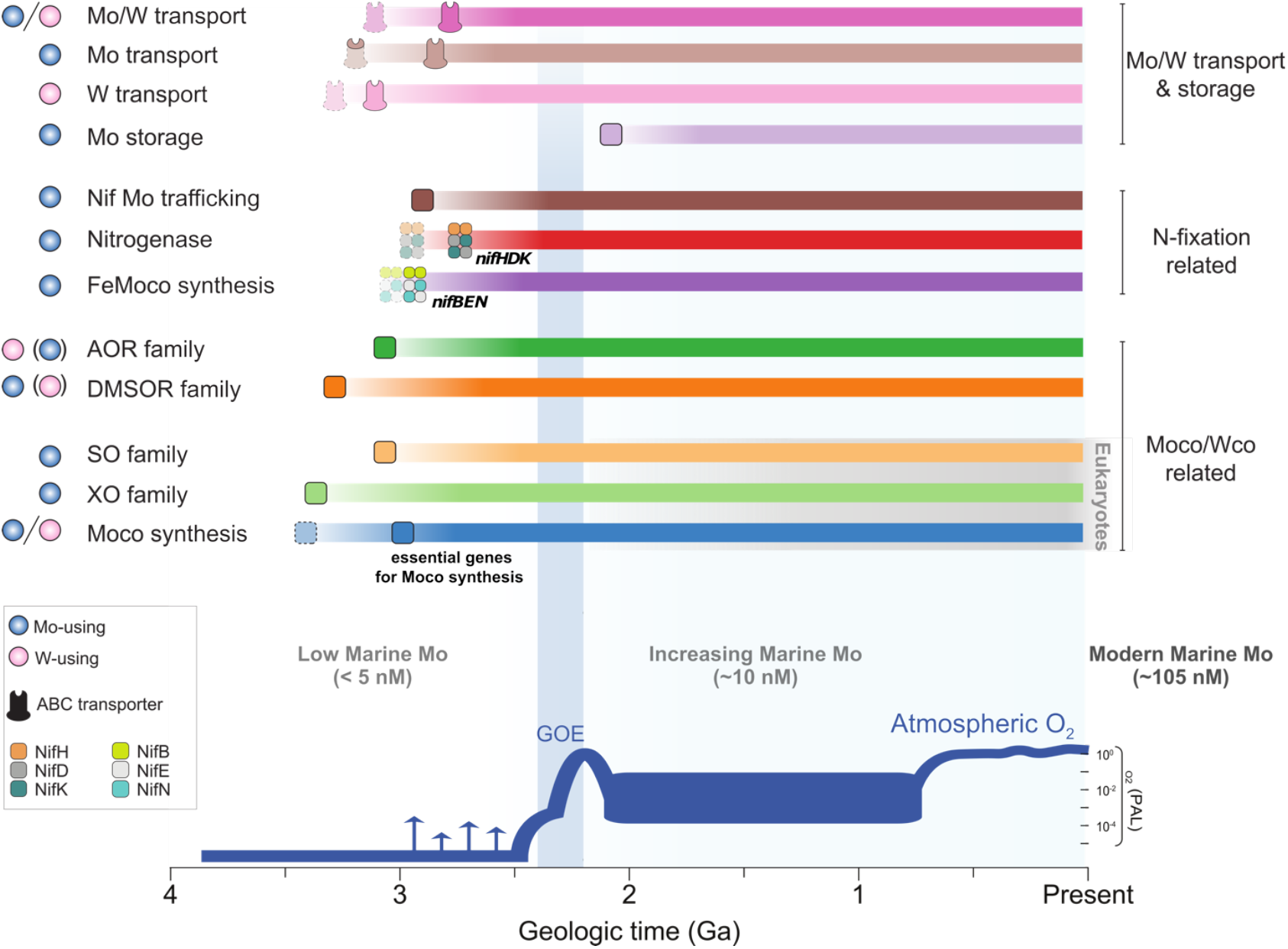
Evolutionary synthesis model for “building” of biological Mo-utilization across Earth history. Shapes at the start of bars correspond to the midpoint date of the earliest gene event recorded for genes within a given group. The gene groups “Mo/W transport”, “Mo transport”, and “W transport” include all three subunits for the respective Mo(/W) ABC-type transporter. The faded transporter shape at the start of bars for these groups corresponds to the midpoint date of the earliest gene event recorded among all three transporter subunits (i.e., the earliest phylogenetic evidence of a single transporter subunit existing). The second filled-in transporter shape corresponds to the earliest phylogenetic evidence of a full ABC-type transporter existing. For the “Mo transport” group, the substrate-binding subunit of the ABC-type Mo transporter (ModA) experienced the earliest gene event among all three subunits (based on the midpoint date) and is filled-in in the transporter shape to denote this. For the “FeMoco Synthesis” and “Nitrogenase” groups, shapes corresponding to three genes essential for FeMoco synthesis (*nifBEN*) and nitrogenase activity (*nifHDK*) are shown. The first faded shape corresponds to the earliest lineage experiencing a gene event among each respective set of genes. The second filled-in shape corresponds to the earliest phylogenetic evidence of all three essential genes existing for the respective group. For the “Moco Synthesis” group, the first faded box corresponds to the earliest lineage experience a gene event among all Moco synthesis genes. The second filled-in box corresponds to the earliest phylogenetic evidence of 9 essential Moco biosynthesis genes existing. Midpoint dates were taken from reconciliation results using the CIR clock model with an HGT cost of 3 (default in ecceTERA) and should be interpreted as estimates for the earliest phylogenetic evidence of Mo/W-related genes per category. For earliest gene events that were speciation events, the right node date was used instead of midpoint date as speciation

Within each category, individual Mo/W proteins form distinct clusters, including the Moco/Wco-dependent enzyme families (XO, DMSOR, SO, AOR), nitrogenase proteins, and different subunits of Mo/W transporters. Distinct clusters also emerge among certain Moco synthesis proteins, reflecting both shared catalytic roles – such as transferase activity – and evolutionary histories. Many Mo/W-related proteins have identifiable homologs that are not Mo/W-related (Figure S4),offering a window into their evolutionary origins. For example, the molybdate binding A subunit of the Mo transporter (ModA) is a member of a broader clade of substrate-binding proteins that includes those specific to compounds like SO_4_^2-^ and phosphate (e.g., H_2_PO_4_^-^)^31^. This supports the hypothesis that this subunit shares an evolutionary origin with non-Mo related transporters.

### Estimated timing for the rise and diversification of Mo proteins

Our analyses have characterized the modern distribution and occurrence patterns of Mo/W-related enzymes, outlining their potential evolutionary history. To contextualize these analyses within geological time, we performed gene tree-species tree reconciliations to estimate the timing of the rise and proliferation of Mo-/W-related proteins. Chronograms from a previous study^26^ were used for this analysis. Briefly, the species tree was anchored in time using eight fossil calibration points and one of three clock models, namely Cox-Ingersoll-Ross (cir), lognormal (ln) or uncorrelated gamma multipliers (ugam), resulting in three different age-calibrated trees (see methods for details). Below, we report dates that consider results from all three clock models tested.

Our results suggest that the utilization of Mo/W in enzymes extends back to the Paleo/Mesoarchean (Figure 3). Estimates of the earliest gene events associated with Mo/W enzymes date back to ~3.7-3.4 Ga depending on the clock model with the LN model predicting the earliest estimates and the UGAM model predicting the most recent estimates (Figures S5-S12). Enzymes associated with the earliest gene events include those involved in the core Moco synthesis pathway, for which a full synthetic pathway likely existed by the Mesoarchean (~3.0-2.7 Ga). Similar evidence suggests the existence of a full W transport system (TupABC) dating back to ~3.1-2.0 Ga, while a full Mo transport system (ModABC and/or WtpABC) is estimated to have emerged around the Mesoarchean to early Paleoproterozoic (~3.0-2.4 Ga).

Among molybdoenzymes, the DMSOR, XO, and AOR families are the most ancient, with their earliest gene events dating back to ~3.5-3.0 Ga. Gene events for these molybdoenzymes remain sparse in the Archean but become increasingly frequent in the Paleoproterozoic. This pattern is consistent with the widespread use of molybdoenzymes across different lineages later in time, potentially in response to the establishment of a fully functional Moco synthesis system. These trends continued with incorporation of Mo into additional enzymes during the Proterozoic, as shown by the more recent gene events of other molybdoenzymes (e.g., torZ, bisC, AOX, XDH). Notably, these later-evolving enzymes are predominantly associated with modern aerobic-respiring organisms and eukaryotes (see Figure S3).

Our findings corroborate previous phylogenomic^4^ and geochemical^5^ studies that date Mo-nitrogenases to an early origin on an anoxic Earth. We also identify maturases (NifB and NifE), which are part of the machinery for FeMoco synthesis in modern organisms, existing as early as the Archean. These proteins share close homology with the catalytic subunits of nitrogenase (NifK and NifD), and phylogenetic studies have proposed that the latter may have emerged from maturase-like predecessors^32^. However, our results cannot discern the evolutionary sequence of events for *nifEB* and *nifDK* genes. Instead, they only indicate that all four genes share ancient origins tied to anoxic conditions on the early Earth. Overall, the ancient emergence of molybdoenzymes across different enzyme families suggests that Mo and W were used by early life to catalyze diverse biochemical pathways. This underscores the lasting importance of these elements, persisting from the Paleoarchean to the present. events correspond to a split in a lineage containing a given gene. Events occurring on terminal branches were not considered when determining earliest events. Note that for the “Moco synthesis” and “AOR family” groups, earliest events from the random reconciliation results determined by ecceTERA were used because the genes with the earliest gene events in these groups included a large number of sequences (see methods for further discussion). Results from the symmetric median reconciliation were used for all other groups. A plot showing changes in levels of atmospheric O_2_ adapted from Lyons et al. (2014)—see ref. in the methods—is shown at the bottom for reference. Estimates for marine Mo concentrations were derived from Johnson et al. (2021)^12^. The gray shading refers to the divergence of Eukaryotic Moco synthesis, XO family, and SO family genes after the evolution of Eukaryotes. Blue spheres represent Mo usage, while pink spheres represent W usage by proteins within each category.

## DISCUSSION

Our results reveal the widespread use of Mo across modern organisms inhabiting a variety of environments and suggest that similar broad-scale Mo-usage patterns were already established in early organisms dating back to ~3.5-3.0 Ga. This challenges the traditional view that Mo availability significantly constrained early biological processes on an anoxic Earth.

Reconciliation results indicate that some of the earliest Mo-/W-related proteins to evolve were involved in Moco biosynthesis. The ancient origin of this biosynthesis pathway is well-supported by the ubiquitous distribution of its proteins across the current tree of life and their conservation among all domains of life^24,27^. Indeed, the same key enzymes forming the prokaryote Moco/Wco-biosynthesis pathway have close eukaryotic homologs which catalyze the same fundamental reactions. The core enzymes that constitute the modern Moco synthesis pathway trace their origins back to the Mesoarchean, suggesting that a complete Moco synthesis pathway may have existed at this time (Figure 3). This coincides with the emergence of molybdoenzymes from different families and Mo(/W)-transport systems. Collectively, these results may indicate that ancient organisms were already investing in Mo(/W) to use for cellular functions on an anoxic Earth.

The geologic record of black shales has been interpreted as evidence of low Mo concentrations in Archean seawater, raising the question of how distinct Mo-enzyme families emerged during this time. One potential explanation addressing this “Mo paradox” may be that suitability to biological functions was a driving factor for choosing Mo. Mo, when incorporated into enzymes, can catalyze reactions spanning a wide range of redox conditions and substrates^1,33^. Consistent with this idea, our analyses support an ancient phylogenetic origin for Mo-utilizing enzymes involved in a variety of essential metabolisms important to C, N, and S cycling (Figures S5-12). Therefore, the antiquity of several distinct Mo-enzyme families suggests that early life could have capitalized on the redox flexibility of Moco, enabling access to a wide-range of substrates and conferring a selective advantage to those capable of using Mo in metabolic reactions. We propose that the functional versatility of Mo-based enzymes has persisted through deep time, reflected in today’s widespread phylogenetic and environmental distribution of molybdoenzymes across different temperatures and pH conditions (Figure 2). Thus, Mo likely played a pivotal role in the earliest energy-harnessing reactions that fueled life long before the GOE. Throughout the Archean, Mo could have been sourced from submarine hydrothermal vents^34,35^ and released into seawater as either dissolved molybdate^36^ or molybdate sorbed onto iron sulfide nanoparticles^37^. The widespread utilization of Mo during the Archean suggests that marine Mo were present in higher concentrations than previously thought^38^, thereby providing a crucial driver for the early emergence of Mo-enzymes.

Intriguingly, the emergence and diversification of molybdoenzymes around the GOE (Figures S5-12), when oxidative weathering began delivering increased amounts of Mo from land to the ocean, suggests that Mo-based biochemistry could have evolved in response to a shift in Mo supply, i.e., from hydrothermal vents^38^ to riverine input. Indeed, many of these molybdoenzymes are associated with aerobe-associated genomes, further supporting the impact of environmental redox conditions on nutrient pathways, such as S cycling (DmsA, SoxC, SoeA), NO_3_^-^ reduction (NasC, NasA), or methane metabolism (TorZ) (Figures S5-12). Oxygenation may have facilitated the evolution of new metabolisms taking advantage of increasingly available oxidized species, with Mo serving as the preferred catalyst.

Mo storage proteins (MosAB) were the only proteins in our analysis whose earliest gene events always post-date the Archean regardless of which clock model is applied (~2.1-1.4 Ga). This presents a curious case as to why these proteins would have emerged during a time of relatively higher predicted marine Mo^13^ (though likely still lower than in the modern oceans^39^). A recent study suggested that competition for Mo between N_2_-fixers and NO_3_^-^ reducers, both relying on molybdoenzyme-catalyzed metabolisms, generated a selection pressure following the GOE that drove molecular innovations tied to Mo-usage^40^. The relatively late evolution of MosAB storage proteins may reflect a Mo-conserving strategy developed by organisms facing Mo scarcity exacerbated by intensified competition for Mo following the GOE. Of note, the distribution of MosAB in the tree of life is restricted to select taxa (Figure 2). This may reflect a later potential advantage of Mo storage for organisms inhabiting specific low-Mo environments^41^ such as highly productive euxinic waters due to increased SO_4_^2-^ availability after the advent of oxidative weathering of crustal pyrite^42^. Alternatively, Mo storage proteins could have been broadly lost in most organisms following the concomitant increase in seawater Mo concentrations driven by higher O_2_ levels.

Alternate theories addressing the Mo paradox lean towards W-dependent metabolisms preceding Mo-dependent ones^43^. Similar to Mo, W can bind pyranopterin to form Wco, which is structurally analogous to Moco. In modern oxygenated marine systems, Mo is significantly more abundant than W, but this disparity decreases under sulfidic and anoxic conditions^43^. Given their similar chemistry^14^ as well as the reported interchangeability between Mo and W in certain metalloenzymes^16^, and their inverse trends in response to O_2_^43^, some theories propose that the earliest molybdoenzymes initially utilized W before incorporating Mo^43^. Our reconciliation data indicate that some of the earliest evolving enzymes include obligate W-dependent AOR as well as facultative W-dependent Fmd/Fwd and Fdh (both of which have known Wco-incorporating and Moco-incorporating homologs)^29,30^. Nonetheless, our data also support the Archean origins of other Mo-specific enzymes, such as xanthine dehydrogenase and sulfite oxidase. This demonstrates the concurrent importance of both Mo and W for early life, with evidence of Mo/W-transport systems (e.g., Wtp, Mod, and Tup) evolving between the Archean and early Paleoproterozoic. The distinct ancestries and limited sequence identity between Mod and Tup transporters, combined with their high ligand-specificities for Mo and W^20^, respectively, support the hypothesis that these transport systems evolved independently to meet the specific metal needs of early life, strongly suggesting sufficient Mo bioavailability to drive the evolution of Mo-specific enzymes in the early Archean^38^.

The working conditions of molybdo-/tungstoenzymes, particularly in relation to temperature, differ^33^, as reflected in the ecological separation of W- and Mo-utilizing organisms in contemporary environments (Figure S2). On the chemical level, tungstoenzymes require a higher temperature to attain optimal catalytic activity, which may explain their ubiquity in contemporary thermophilic archaea^33^. In addition, W-based redox reactions generally operate at relatively lower redox potentials and over a narrower range than molybdoenzymes^33^. Combined with evidence that W can antagonize molybdoenzymes and vice versa^44^, it is possible that Mo- and W-based enzymes evolved concurrently rather than sequentially (i.e., W before Mo), allowing early organisms to catalyze different biochemical reactions. In fact, the ability to use both transition metals may have even provided a selective advantage for organisms living in systems limited in either metal^45^.

Taken together, our analysis suggests that Mo utilization in biological systems originated during the Paleo-to Mesoarchean, with Moco biosynthesis and molybdoenzymes playing essential roles in early metabolisms, potentially due to the redox flexibility of the latter. Following the GOE, Mo-dependent enzymes diversified into aerobe-associated metabolisms. Competition for Mo drove adaptive strategies such as Mo storage proteins, with their limited distribution reflecting evolutionary trade-offs linked to environmental Mo availability. Most Mo/W-related proteins investigated in this study have their earliest gene events estimated to date back to the Archean and the period coinciding with the GOE. These enzymes catalyze diverse pathways, underscoring the critical role of Mo and W in early biological productivity and (more broadly) nutrient cycling on the early Earth. The relative richness of Mo prior to the oxidation of Earth’s atmosphere and ocean^36,46^, combined with these catalytic functions, suggests that the anoxic early Earth supported greater Mo-based biochemical versatility than previously recognized.

## ACKNOWLEDGEMENTS

This work was supported by the National Aeronautics and Space Administration (NASA) Interdisciplinary Consortium for Astrobiology Research: Metal Utilization and Selection Across Eons, MUSE [80NSSC17K0296] (BK) with additional support from the Paglia Post-Baccalaureate Research Fellowship from Carleton College (ASK) and the NASA Postdoctoral Program, administered by Oak Ridge Associated Universities under contract with NASA (MSS). Authors also acknowledge additional funding support from a NASA Astrobiology Program Grant [80NSSC18KO829] (RA) and a NERC Frontiers grant (NE/V010824/1) (JSB and EES). We thank Evrim Fer and Amanda Garcia for their assistance in data visualization; Holly Rucker, Kaustubh Amritkar, Ariel Anbar, Lance Seefeldt, and the members of the MUSE ICAR for valuable feedback and discussions; the Center for High Throughput Computing (CHTC) at the University of Wisconsin–Madison for providing computing resources; and the University of Wisconsin–Madison Department of Bacteriology Bioinformatics Core for the input.

## AUTHOR CONTRIBUTIONS

ASK: Study design, data acquisition and analysis, writing (original draft), editing; MSS: data acquisition and analysis, writing-editing; JSB: writing-editing, provision of molecular clocks and advice on reconciliation analyses; ES: editing, REA: editing, provision of molecular clocks and advice on reconciliation analyses, KK: writing-editing, BK: Conceptualization, study design, data analysis, writing (original draft), editing, supervision, funding.

## DATA AND CODE AVAILABILITY

The data used for all analyses presented in this article along with supporting code for data analysis can be found at https://github.com/kacarlab/MoW_evo.

