## Supplemental Information for "Biological molybdenum usage stems back to 3.4 billion years ago"

### MATERIALS & METHODS

#### Genome and protein dataset curation for phylogenetic distribution of Mo-genes

Genome and protein sequence data were curated from 1,609 bacterial, archaeal, and eukaryotic organisms. The selected organisms are a subset of those included in a previously constructed reference tree of life[1]. Organisms were selected based on the availability of protein sequence data associated with each genome assembly (downloaded from NCBI[2], August 2023). In other words, genomes of taxa with no associated protein sequence data available on NCBI were excluded from our analyses. As a final step, all bacterial and archaeal genomes including MAGs and isolate genomes were filtered for quality (completion ≥ 90%, contamination ≤ 5%) of associated genome assemblies, predicted by CheckM2[3] using default settings.

#### Protein functional annotation

Nitrogenase (Nif/Vnf/AnfHDGK) subunit proteins from the curated genome dataset were manually annotated with KEGG Orthology identifiers ("KO numbers") as follows. Candidate nitrogenase sequences were identified by HMMER v.3.3.2 (hmmsearch) using KOfam HMM profile[4] queries (downloaded July 2023) against a local database of all protein sequences associated with the collected genome assemblies (NifH: K02588, NifD: K02586, NifK: K02591, VnfG: K22898). hmmsearch was performed with full-sequence (-E) and domain (--domE) E-value thresholds of < $1 \times 10^{-5}$. Target sequences from each hmmsearch run were aligned to previously published nitrogenase protein sequence datasets[5,6] by MAFFT v.7.490[7] (alignment strategy was automatically selected by the software using the –auto flag) and used to build maximum likelihood phylogenetic trees by FastTree v.2.1.11[8] (WAG model) with 1000 bootstraps. The WAG model was chosen from one of two available models (WAG or JTT) in the Geneious FastTree plug-in software, where WAG has been shown to outperform JTT[9]. Manual annotations of nitrogenase sequences were made based on phylogenetic clustering, using previously described nitrogenase protein clades as reference[5,6,10], as well as the presence of known, nitrogenase metallocluster ligands (e.g., FeMo-cofactor His442 and Cys275 ligands; site index from *Azotobacter vinelandii* NifD).

All other sequences in the protein dataset were annotated with KO numbers by HMMER v.3.3.2 (hmmscan; http://hmmer.org/) against a local copy of the KOfam HMM profile database[4] (downloaded July 2023) using the University of Wisconsin-Madison Center for High Throughput Computing (CHTC) servers. hmmscan was performed with full-sequence (-E) and domain (--domE) E-value thresholds of < 1 x $10^{-5}$. Each protein query was annotated with the KO number target with the lowest full-sequence E-value. It should be noted that KEGG does not distinguish between the Mo-using and W-using orthologs of formylmethanofuran dehydrogenase. Thus, all analyses on the subunit harboring the Mo/W-containing active site (FwdB/FmdB: K00201) of these enzymes reflect combined results for both genes.

### Identification of Mo-related proteins

Initial Mo-related protein targets were identified by a literature search[11–15]. Proteins annotated as described above were grouped by corresponding KO numbers, aligned by MAFFT[7] and used to generate an HMM profile database by hhmake in HH-Suite[8]. HMM profiles corresponding to Mo-related gene targets identified by the literature search were used as queries to perform HMM-HMM homology search against the generated HMM profile database using hhsearch in HH-Suite. HMM profiles with detected homology to the initial Mo-related HMM profile queries were manually inspected for the presence of Mo-related functional domains, based on Mo-related pfam annotations available for KEGG Orthologs in the KEGG database. Confirmed homologs were used to expand the Mo-related protein target dataset. The final Mo-related protein dataset contained 102 unique KO number targets.

### Mo-related protein network

The HMM-HMM homology search output was used to build a protein homology network using the NetworkX python library. Edges were drawn with a homology threshold of log-eval > 15.

### Gene count matrix

Mo-related gene counts were mapped onto a previously constructed Hug et al. reference species tree of life[1]. R packages Ape v.5.7-1[16] and ggtree v.3.8.0[17] were used to prune taxa absent from the curated genome dataset and tree visualization, respectively. This resulted in a set of 865 genomes present in the time-calibrated tree. Mo-related gene counts per genome were generated and visualized using a KO number annotation E-value threshold of 1e-10, determined by hmmscan as described above.

### Gene co-occurrence analysis

Pairwise Spearman's rank correlation coefficients were analyzed across presence/absence data of Mo-/W-related genes, and visualized as a hierarchically clustered matrix (using the parameter order = 'hclust') with the corrplot package[18] in R.

### Ecological attribute prediction and correlation analysis

Optimal growth temperature, pH, habitat, and $O_2$ tolerance metadata were predicted for selected genomes from Hug. et al. (2016)[1] as follows. The *Oxyphen* package[19] was used to predict oxygen metabolism. The *OGT_prediction* tool[20] was used to estimate optimal growth temperature with the bacterial and archaeal regression model that excluded information from the 16S rRNA gene as well as genome size (*superkingdom-all_species-all_features_ex_genome_size_and_rRNA.txt*). Predictions for pH were made using the Ramoneda et al (2023)[21] model. Habitat preference and richness were determined with ProkAtlas[22] using a 16S nucleotide identity of 97% and a sequence coverage threshold of 150 base pairs. Spearman's rank correlation coefficient between environmental trait metadata and number of Mo/W genes were calculated in R.

### Age estimation of Mo-/W-related proteins

To ascertain the timing of gene duplication, loss, horizontal gene transfer (HGT), and speciation events for Mo-/W-related genes, we reconciled phylogenetic trees for each gene of interest against a molecular clock containing 865 genomes selected to represent the full diversity of life on Earth (first described in Boden et al. Nat Commun 2024[23]). For each Mo-/W-related gene of interest, we identified genes in the 865 genomes included in the molecular clock via their KO number annotations as described above. Because KEGG does not distinguish between *vnf* and *anf* (alternative nitrogenase) sequences, these genes were excluded from the reconciliation analyses. Sequences were subsequently grouped by KO number annotations. KO number groups with fewer than 10 sequences were combined with the closest KO number homolog determined by HMM-HMM homology searches (described above), yielding 76 different groups.

Protein sequences in each KO number group were aligned in MAFFT[7] v7.505 with E-INS-i and trimmed to remove column positions comprising >85% gaps with trimAl v1.4.rev15[24](-gt 0.15). Trimmed sequence alignments were then used to construct phylogenetic trees by IQ-TREE v.2.1.4[25] with 1000 ultrafast bootstraps and the best-fit substitution model chosen by ModelFinder[26]. The resulting gene trees were reconciled with four previously constructed age-calibrated species trees (from Boden et al. Nat Commun 2024[23]) using ecceTERA reconciliation software[27] to estimate the timing of gene events. Further details on the construction of chronograms are described in Boden et al. Nat Commun. 2024[23]. Briefly, a species tree was constructed using 16 different ribosomal proteins from 865 lineages. This tree was anchored in time using PhyloBayes[28] with 8 fossil calibration points and a uniform distribution on the root prior and one of three clock models, namely Cox-Ingersoll-Ross (cir), lognormal (ln) or uncorrelated gamma multipliers (ugam). This resulted in three age-calibrated species trees, each made with a different clock model (because there is no current consensus on which is most appropriate for estimating ages in deep time). An additional age-calibrated species tree was made with the ln clock model by varying one of the eight calibration points to enforce an earlier origin of methanogenesis at > 3.46 Ga instead of > 2.7 Ga in-line with recent findings[29–31]). This served as a sensitivity test to investigate the effect of changing this calibration point.

Default settings were implemented in ecceTERA for reconciliations, specifying amalgamation of gene trees with the amalgamate=true parameter and allowing transfers from extinct branches (compute.TD=true). As previously reported[23], the default costs for gene events assume that HGT is more “expensive" than speciation, duplication, and loss to reduce genome size variation between parent and daughter lineages (David & Alm, 2011[32]). Consistent with previous methods[23], we also report reconciliation results using different HGT costs (2, 4, and 6, alongside the default of 3, Figure S20) to account for scenarios involving greater genome size variation. ecceTERA output was parsed using the recon2xml.pl Perl script provided by ecceTERA software and a set of custom python scripts (several adapted from Mateos et al Sci Adv 2023[33]). Reconciliation data associating gene events with dates corresponding to nodes of the species tree were visualized in R (Figures S5-S12).

Several of our gene trees (MOCS1, moaA/CNX2; ygeS/xdhA; GPHN, moeA; mobA; moaC/CNX3; aor; moaD/cysO; sreA, phsA/psrA, padB, frdC; moeB) had too many sequences for computation of the symmetric median reconciliations by ecceTERA. Thus, for these genes, we report results from the random reconciliation chosen by ecceTERA. Reconciliation output from both the symmetric median reconciliations (when available) and random reconciliations using all clock models are shown in Figures S5-S12.

Caution is warranted when interpreting estimates from tree reconciliations. For example, a gene event can occur anywhere along a branch of the species tree, resulting in a wide range of possible ages. Here, we report midpoint dates (calculated as the average of dates corresponding to the left and right nodes of a branch) for HGT, duplication, and loss events. Speciation events occur when a lineage containing a gene diverges into two species. Thus, dates corresponding to these internal nodes are reported, consistent with previous analyses[23]. Interestingly, we find that older events tend to be more temporally constrained and occur on shorter branches in the species tree compared to more recent events, as previously reported[33]. Plots depicting the full temporal range of gene events (spanning the range between left and right node dates) are available in Figures S13-S19. The prediction of events also heavily relies on the topology of both the chronogram and gene tree being used, meaning that new gene sequence discoveries may alter our interpretations. Consequently, we report results from all three chronogram models tested, in the main text. Figure 3 in the main text depicts a summary of the oldest gene events among groups of analyzed genes, plotted alongside a separate plot adapted from Lyons et al. (2014)[34] depicting changes in levels of atmospheric $O_2$ over time.

Finally, it should be reiterated that our record of the history of ancient genes comes from inferences based on modern genomes. These proteins may have originally bound different metals or performed alternate function in the past compared to their present-day counterparts. In light of these limitations, we focused on broad-scale trends and the relative timing for the emergence and proliferation of genes across the tree of life, while remaining mindful of the possibility that ancient enzymes may have had alternate functions or structures that are no longer observed in extant systems.

**SUPPLEMENTAL FIGURES**

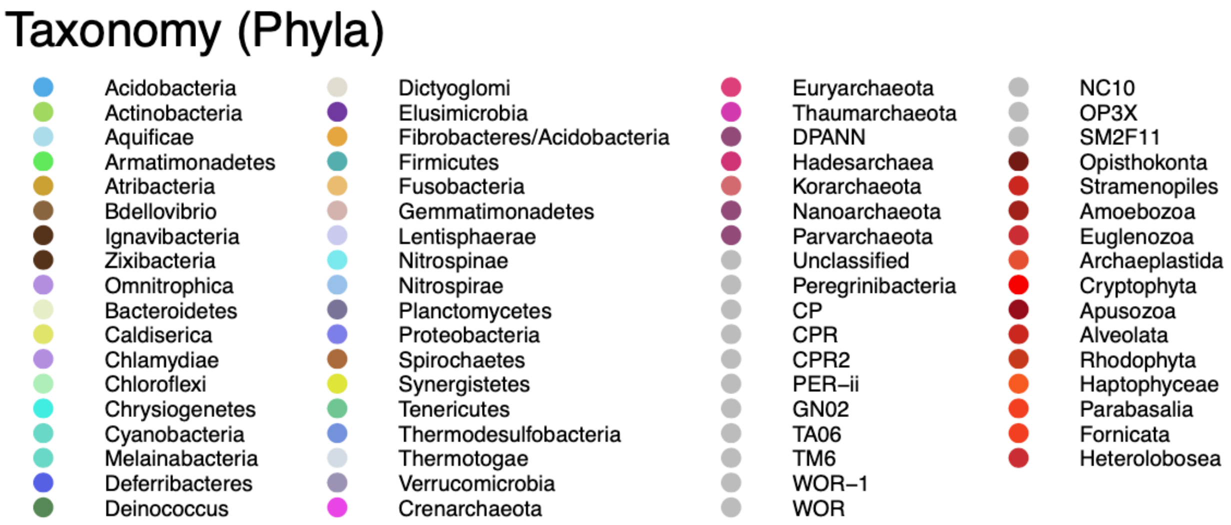


**Figure S1. Legend for the species phylogeny in Figure 1 showing different taxa by color.** Taxa are colored at the level of phylum as presented in Hug et al. (2016) where the phylogeny was derived from. Groups in gray ("Unclassified" through "SM2F11") refer to those that lacked isolated representatives at the time of that study.

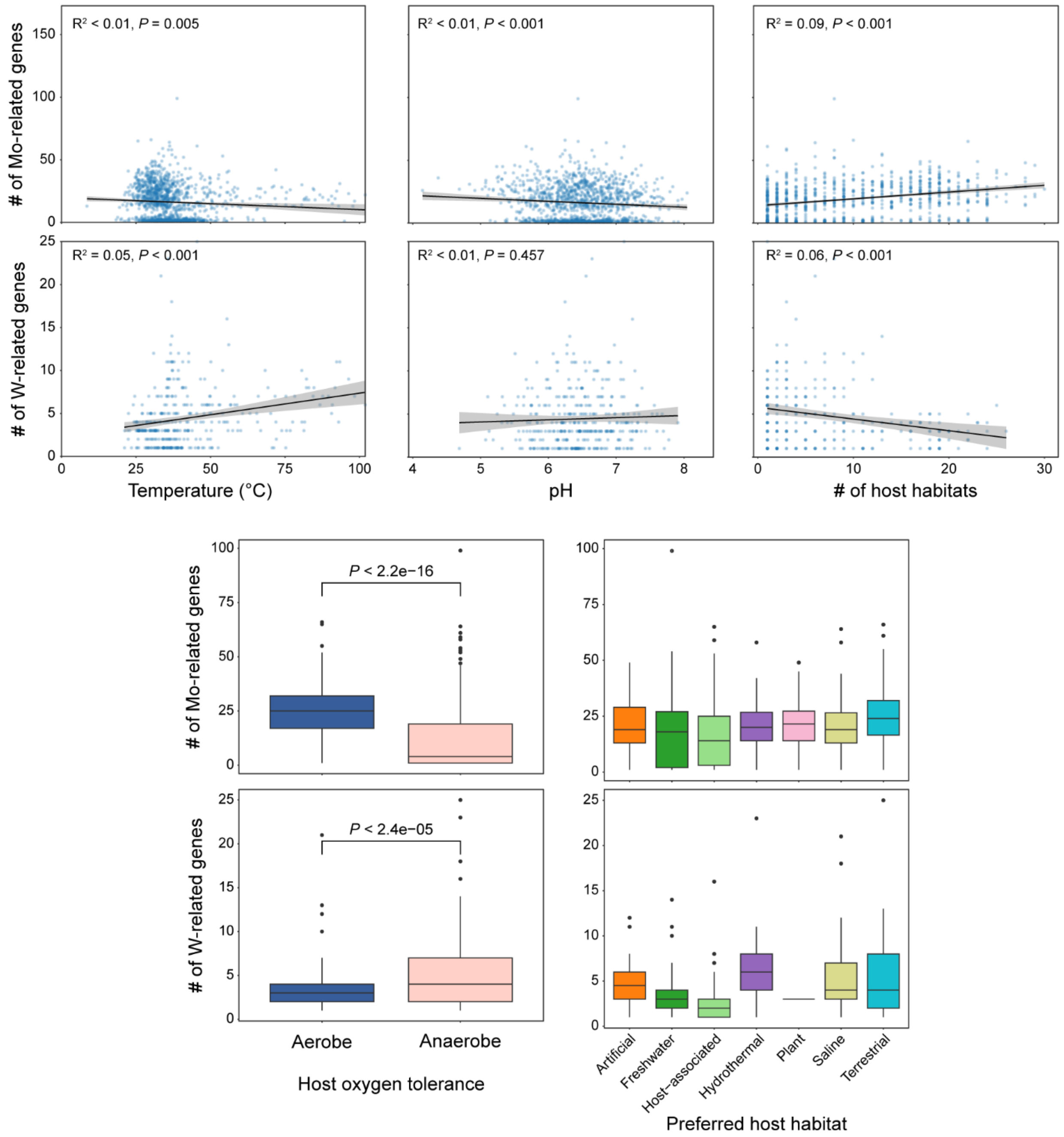


**Figure S2. Correlations between number of Mo/W-related genes and host ecological attributes for analyzed genomes.** Mo-related genes include enzymes belonging to the DMSOR, SO, and XO families, Moco and FeMoco synthesis genes, and genes involved in Mo storage, trafficking, and transport (Mod and Wtp transporters). W-related genes include AOR family enzymes, as well as W-transporters (Wtp and Tup). Scatter plots show linear model fit to the data ($R^2$ and *P* values reported at top). The shaded areas represent standard error of the linear model. P values for host oxygen tolerance data were calculated by the Wilcoxon rank sum test. Solid circles above box plots represent outliers. Weak yet statistically significant correlations are shown between total counts of Mo/W genes in individual genomes and host temperature ($R^2 < 0.01$, $P < 0.005$; and $R^2 = 0.05$, $P < 0.001$ for Mo and W genes, respectively), pH ($R^2 < 0.01$, $P < 0.001$, except for W gene counts where $R^2 < 0.01$, $P = 0.457$), and the number of habitats associated with a given genome ($R^2 = 0.09$, $P < 0.001$; and $R^2 = 0.06$, $P < 0.001$ for Mo and W genes, respectively). Additionally, there are significant differences in the total Mo/W gene counts between aerobic and anaerobic organisms (with $P < 2.2 \times 10^{-16}$ and $P < 2.4 \times 10^{-5}$ for Mo and W genes, respectively), where aerobes tend to encode more Mo genes than anaerobes and anaerobes tend to encode more W genes than aerobes. Given the inverse relationship between molybdate and tungstate abundances with $O_2$ levels in modern marine systems[35,36], the observed trends in Mo/W gene counts suggest that metal availability may, in part, influence metal selection in modern organisms. In general, the directionality of the observed correlations diverges between Mo and W gene counts (with organisms with numerous Mo genes associated with more temperate, alkaline, and oxic environments, and organisms with more W genes associated with hotter, acidic, and anoxic environments), suggesting that organisms with high Mo requirements tend to occupy different ecological niches than organisms with high W requirements. Given the weak correlations between Mo/W gene counts and environmental factors, however, the observed trends between organisms with high Mo requirements and those with high W requirements in the occupancy of different niches may be subtle and instead suggest other factors, such as the specific suitability of each metal for key metabolic pathways, may play a more significant role in determining Mo/W utilization in certain ecological niches.

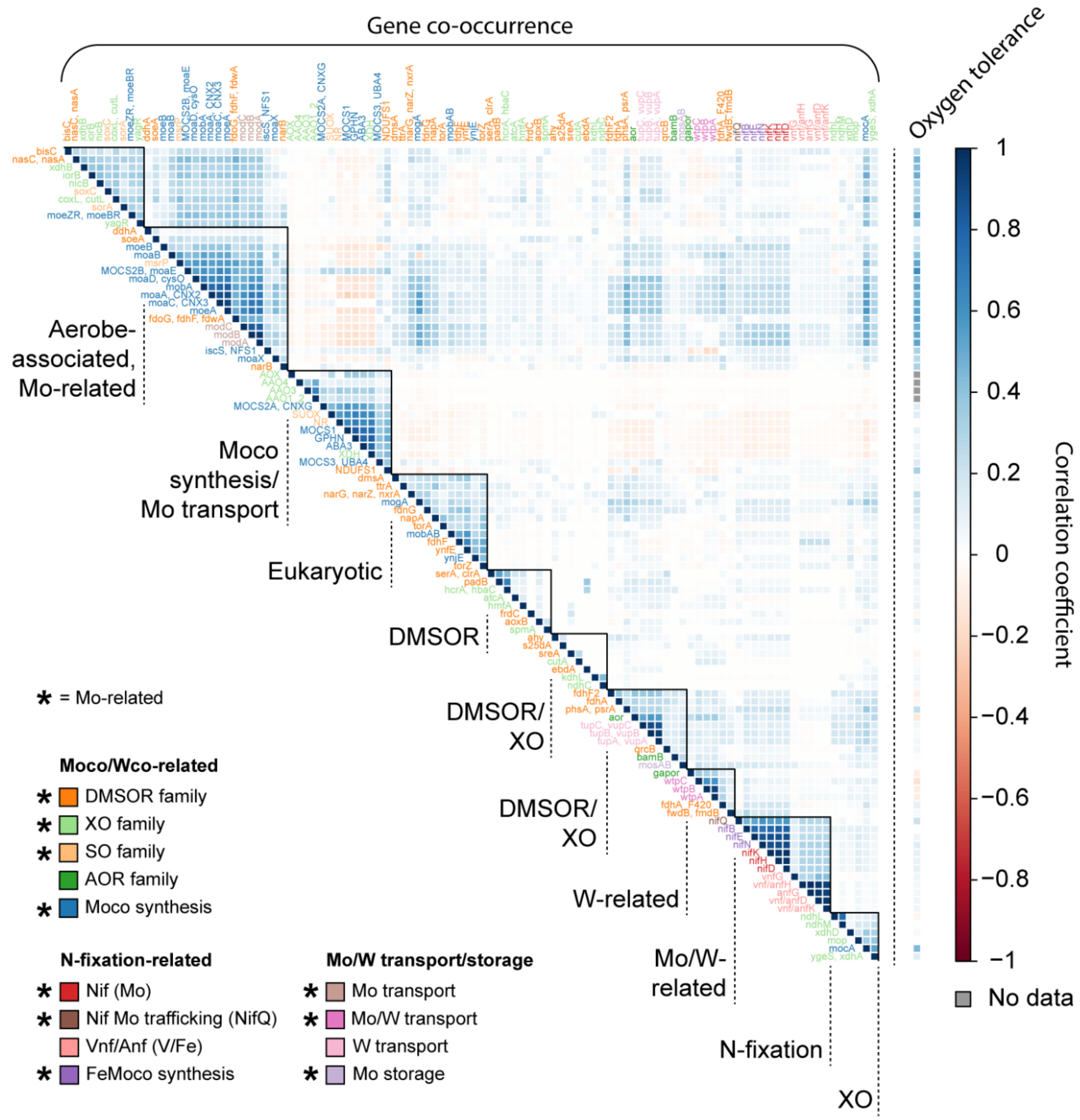


**Figure S3. Genome-level co-occurrence of genes associated with Mo-dependent cellular functions.** The matrix displays Spearman's rank correlation coefficients calculated from binary gene presence annotations for our genome dataset. Groups resulting from hierarchical clustering of the gene correlation data are indicated with bold triangles. Below each group is labeled with notable characteristics and/or the most representative gene categories. The single column on the right displays correlation coefficients between gene presence and oxygen tolerance of the host organism (coded as binary "aerobe" or "anaerobe" annotations), where a positive coefficient indicates a positive correlation between gene presence and an aerobic host lifestyle. Gene presence/oxygen tolerance correlations were calculated only for prokaryotic genomes and were not factored in the hierarchical clustering analysis. Genes with missing gene presence/oxygen tolerance correlation data were not found in prokaryotes and were thus not included in that analysis. Mo-related genes most strongly associated with an aerobic lifestyle tend to co-occur in the same organisms. This includes genes that enable organisms to utilize a wide variety of compounds as sources of N, C, and S, such as nicotinic acid (*nic* genes) for N and C metabolism[37]; and thiosulfate and sulfite (*sox* and *sor* genes) for S metabolism[38]. These expanded metabolic capabilities may reflect a selective advantage in growth and survival for organisms living in oxygenated environments.

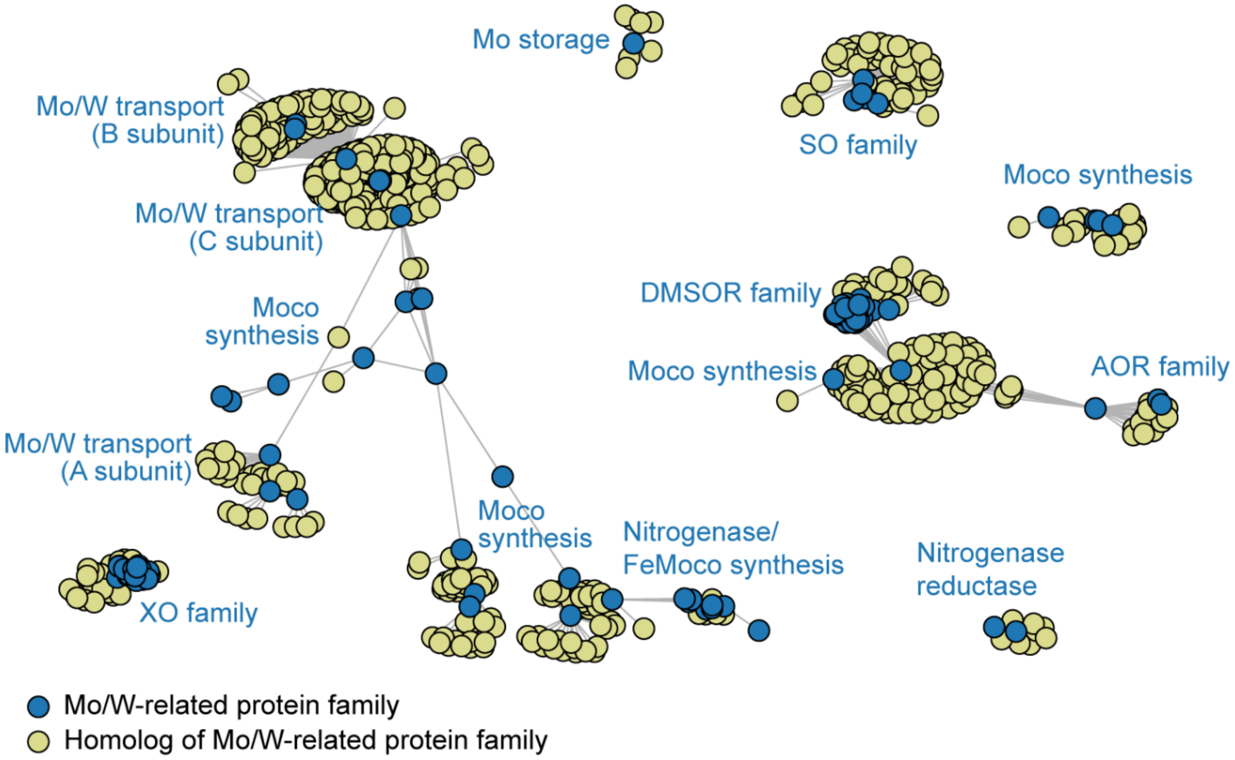


**Figure S4. Similarity network of Mo-related protein families and related homologs.** The network is built from pairwise HMM-HMM comparisons (performed by hhsearch; see Methods and Materials) between Mo/W-related protein families and all other protein families within the KEGG KOFam database. Each node represents an individual protein family and each edge represents detectable homology between protein families ($\log_2$(E-value) < -15 threshold for each HMM-HMM comparison). Labels correspond only to Mo/W-related protein families in each cluster. Blue circles correspond to Mo/W-related protein families, while yellow circles correspond to homologs of these families (see legend).

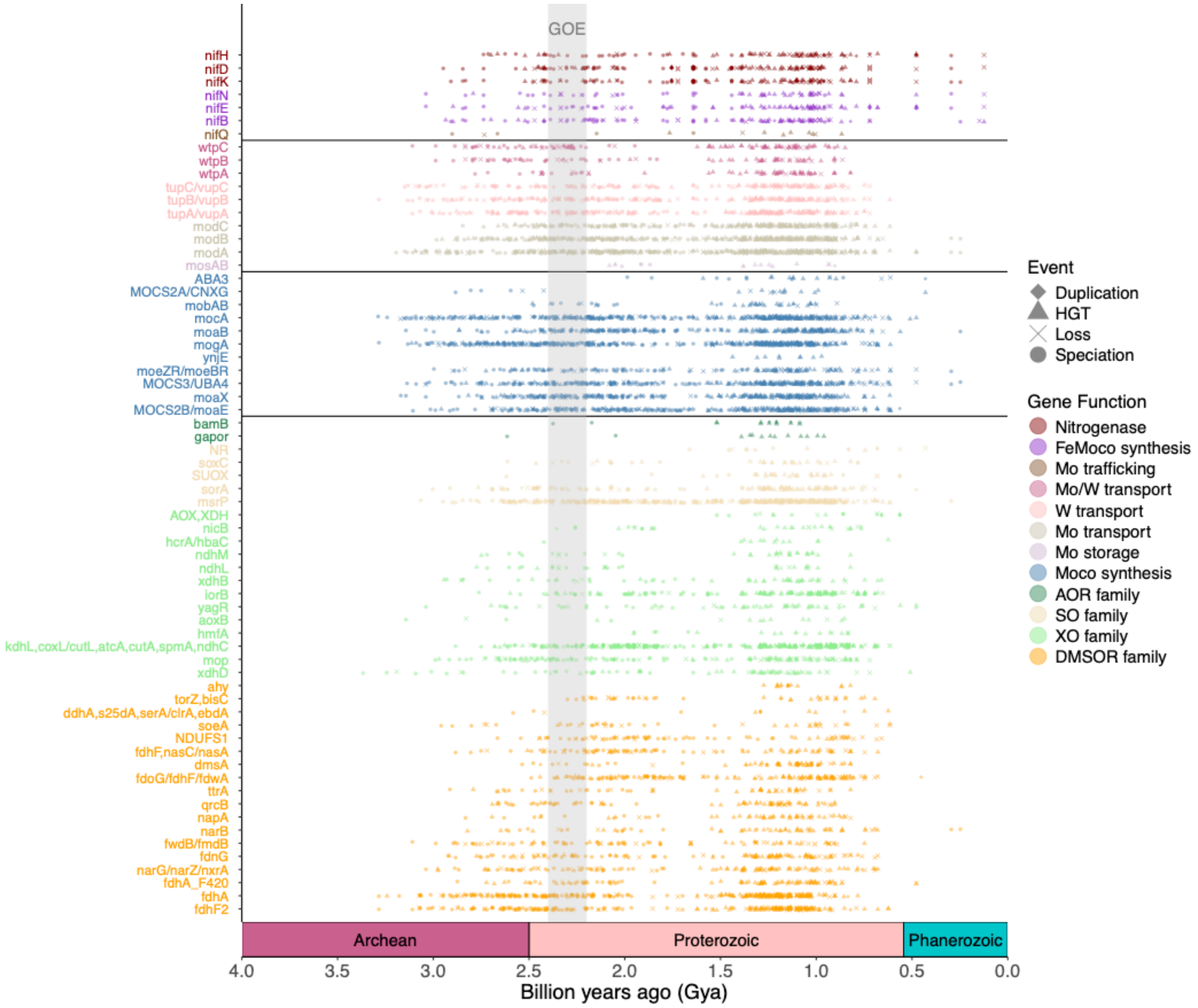


**Figure S5. Gene speciation, duplication, loss, and horizontal gene transfer events for Mo-/W-related genes from the symmetric median reconciliation identified by ecceTERA using the autocorrelated Cox-Ingersoll-Ross (CIR) clock model, plotted across time.** Gene events from the symmetric median reconciliation using the CIR clock model and the ecceTERA default horizontal gene transfer (HGT) cost of 3 are shown above. Each point represents a single gene event where shapes distinguish different gene event types. Only the midpoint date is shown, except for speciation events where the right node date is shown. Events are colored based on gene groups (shown in boxes above). Lines separate the major Mo-enzyme families, enzymes involved in Moco biosynthesis, metal transport/storage proteins, and proteins involved in nitrogenase assembly and function. The gray rectangle marks an estimated time range for the Great Oxidation Event (GOE, ~2.4-2.2 Gya). Note that gene losses (X's) suggest a gene is no longer being used by a lineage and should not be interpreted as evidence of gene usage.

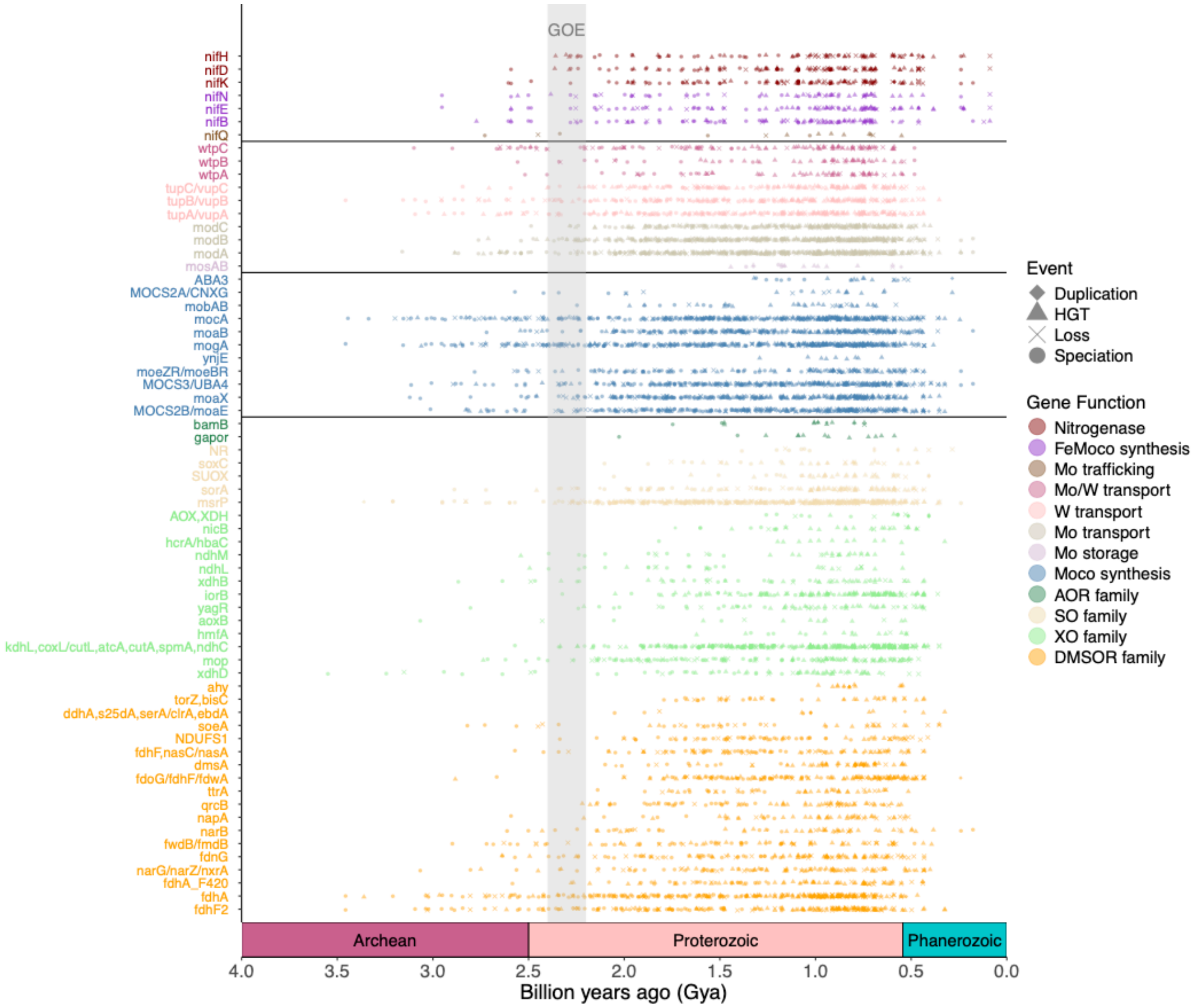


**Figure S6. Gene speciation, duplication, loss, and horizontal gene transfer events for Mo-/W-related genes from the symmetric median reconciliation identified by ecceTERA using the autocorrelated lognormal (LN) clock model, plotted across time.** Gene events from the symmetric median reconciliation using the LN clock model and the ecceTERA default horizontal gene transfer (HGT) cost of 3 are shown above. Each point represents a single gene event where shapes distinguish different gene event types. Only the midpoint date is shown, except for speciation events where the right node date is shown. Events are colored based on gene groups (shown in boxes above). Lines separate the major Mo-enzyme families, enzymes involved in Moco biosynthesis, metal transport/storage proteins, and proteins involved in nitrogenase assembly and function. The gray rectangle marks an estimated time range for the Great Oxidation Event (GOE, ~2.4-2.2 Gya). Note that gene losses (X's) suggest a gene is no longer being used by a lineage and should not be interpreted as evidence of gene usage.

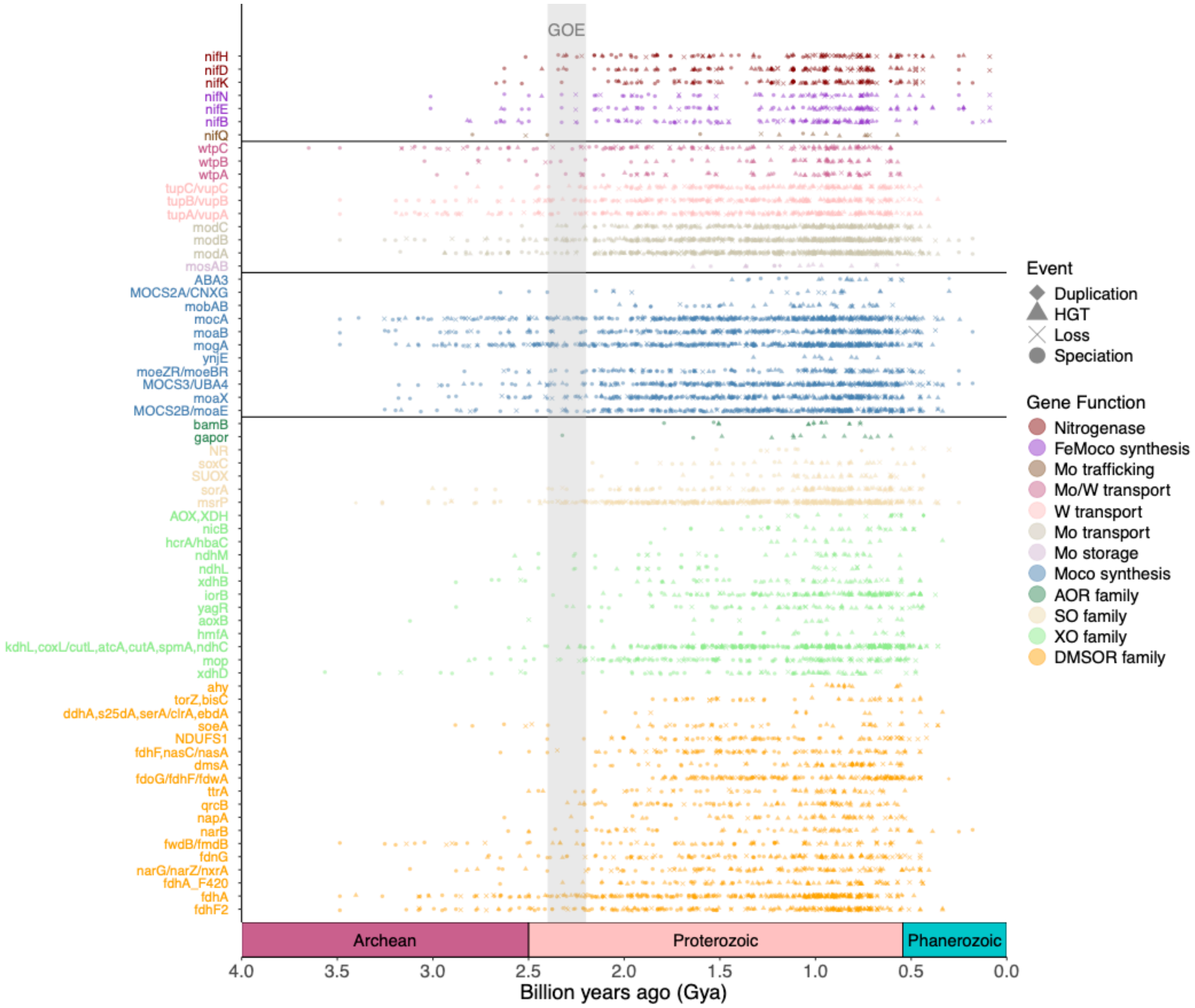


**Figure S7. Gene speciation, duplication, loss, and horizontal gene transfer events for Mo-/W-related genes from the symmetric median reconciliation identified by ecceTERA using a new autocorrelated lognormal (LN) clock model from Boden et al. (2024) generated using an earlier estimation for methanogenesis (> 3.46 Ga), plotted across time.** Gene events from the symmetric median reconciliation using a different LN clock model assuming an earlier estimate for methanogenesis from Boden et al. (2024) and the ecceTERA default horizontal gene transfer (HGT) cost of 3 are shown above. Each point represents a single gene event where shapes distinguish different gene event types. Only the midpoint date is shown, except for speciation events where the right node date is shown. Events are colored based on gene groups (shown in boxes above). Lines separate the major Mo-enzyme families, enzymes involved in Moco biosynthesis, metal transport/storage proteins, and proteins involved in nitrogenase assembly and function. The gray rectangle marks an estimated time range for the Great Oxidation Event (GOE, ~2.4-2.2 Gya). Note that gene losses (X's) suggest a gene is no longer being used by a lineage and should not be interpreted as evidence of gene usage.

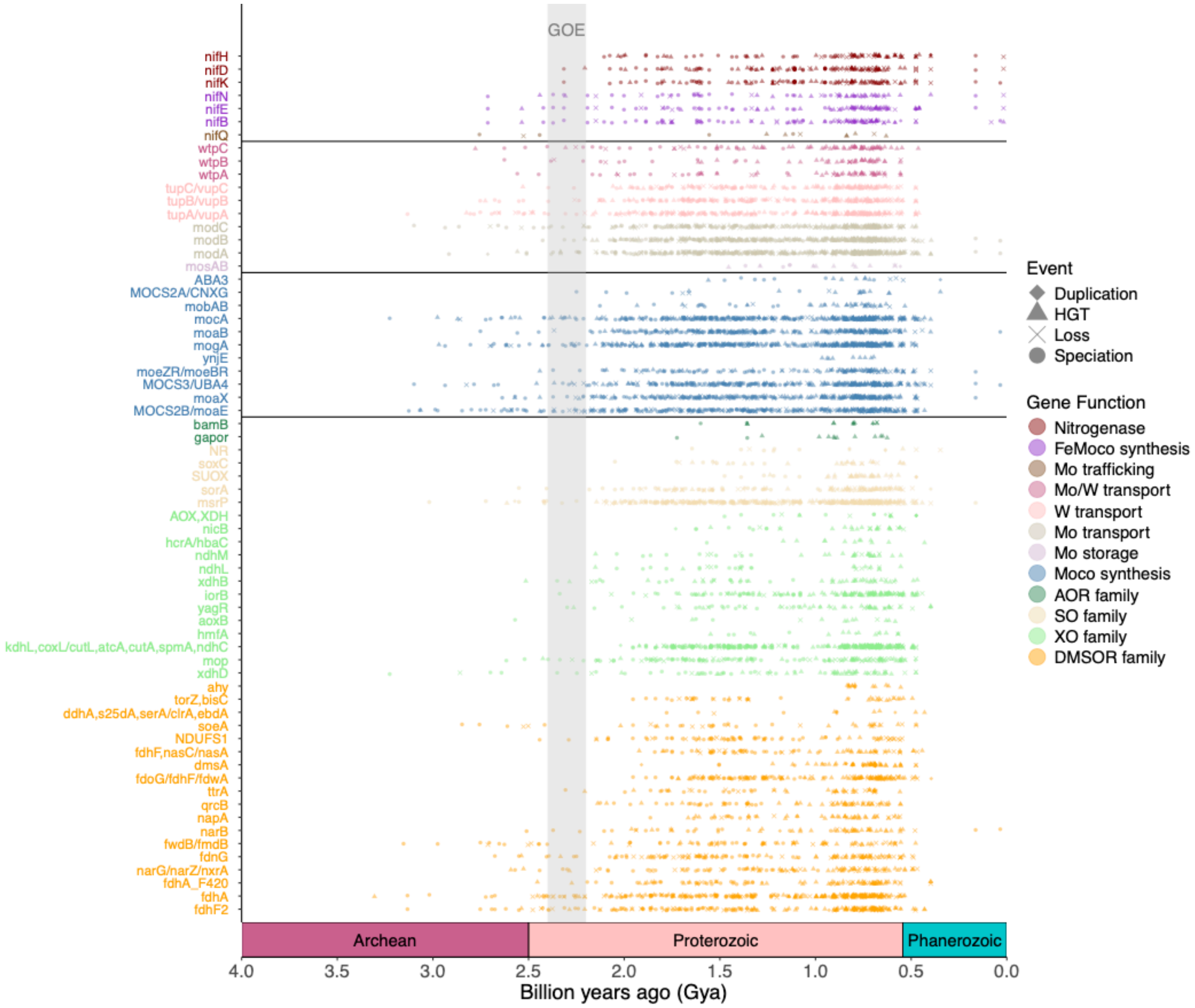


**Figure S8. Gene speciation, duplication, loss, and horizontal gene transfer events for Mo-/W-related genes from the symmetric median reconciliation identified by ecceTERA using the uncorrelated gamma multipliers (UGAM) clock model, plotted across time.** Gene events from the symmetric median reconciliation using the UGAM clock model and the ecceTERA default horizontal gene transfer (HGT) cost of 3 are shown above. Each point represents a single gene event where shapes distinguish different gene event types. Only the midpoint date is shown, except for speciation events where the right node date is shown. Events are colored based on gene groups (shown in boxes above). Lines separate the major Mo-enzyme families, enzymes involved in Moco biosynthesis, metal transport/storage proteins, and proteins involved in nitrogenase assembly and function. The gray rectangle marks an estimated time range for the Great Oxidation Event (GOE, ~2.4-2.2 Gya). Note that gene losses (X's) suggest a gene is no longer being used by a lineage and should not be interpreted as evidence of gene usage.

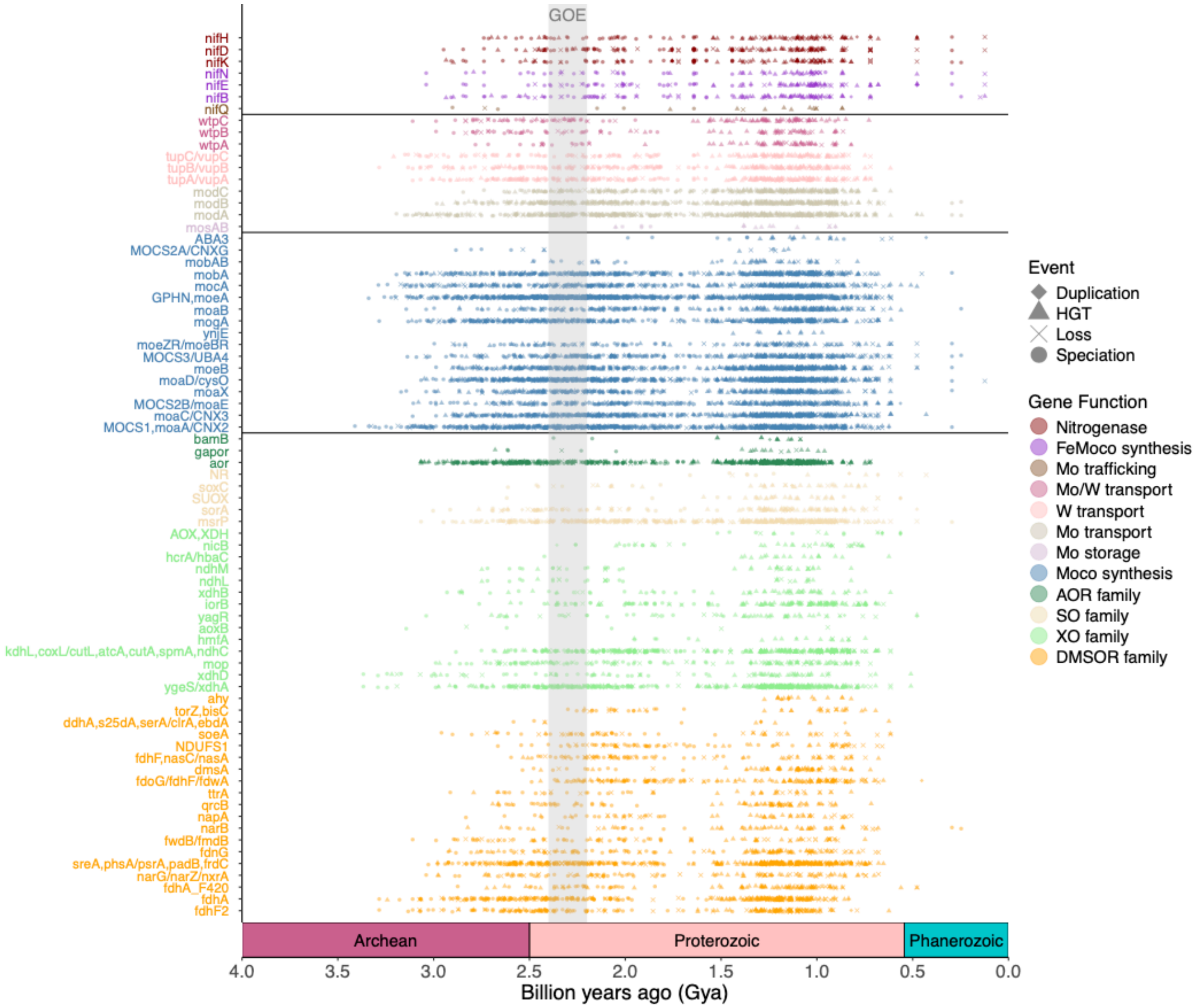


**Figure S9. Gene speciation, duplication, loss, and horizontal gene transfer events from a random reconciliation plotted across time for Mo-/W-related genes as identified by ecceTERA using the autocorrelated Cox-Ingersoll-Ross (CIR) clock model.** Random reconciliation results (as opposed to symmetric or asymmetric median reconciliations) using the CIR clock model and the ecceTERA default horizontal gene transfer (HGT) cost of 3 are shown above. Each point represents a single gene event where shapes distinguish different gene event types. Only the midpoint date is shown, except for speciation events where the right node date is shown. Events are colored based on gene groups (shown in boxes above). Lines separate the major Mo-enzyme families, enzymes involved in Moco biosynthesis, metal transport/storage proteins, and proteins involved in nitrogenase assembly and function. The gray rectangle marks an estimated time range for the Great Oxidation Event (GOE, ~2.4-2.2 Gya). Note that gene losses (X's) suggest a gene is no longer being used by a lineage and should not be interpreted as evidence of gene usage.

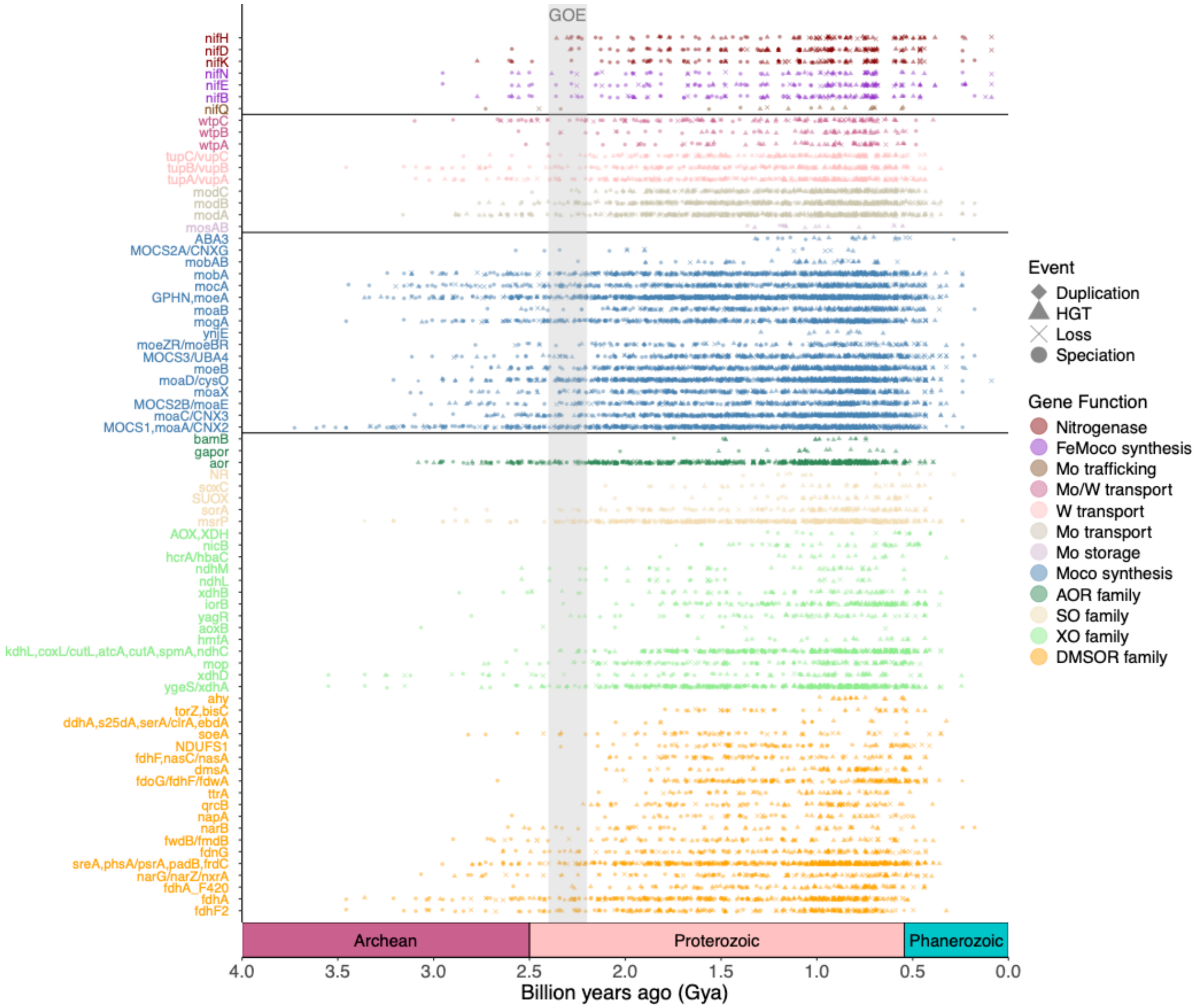


**Figure S10. Gene speciation, duplication, loss, and horizontal gene transfer events from a random reconciliation plotted across time for Mo-/W-related genes as identified by ecceTERA using the autocorrelated lognormal (LN) clock model.** Random reconciliation results (as opposed to symmetric or asymmetric median reconciliations) using the LN clock model and the ecceTERA default horizontal gene transfer (HGT) cost of 3 are shown above. Each point represents a single gene event where shapes distinguish different gene event types. Only the midpoint date is shown, except for speciation events where the right node date is shown. Events are colored based on gene groups (shown in boxes above). Lines separate the major Mo-enzyme families, enzymes involved in Moco biosynthesis, metal transport/storage proteins, and proteins involved in nitrogenase assembly and function. The gray rectangle marks an estimated time range for the Great Oxidation Event (GOE, ~2.4-2.2 Gya). Note that gene losses (X's) suggest a gene is no longer being used by a lineage and should not be interpreted as evidence of gene usage.

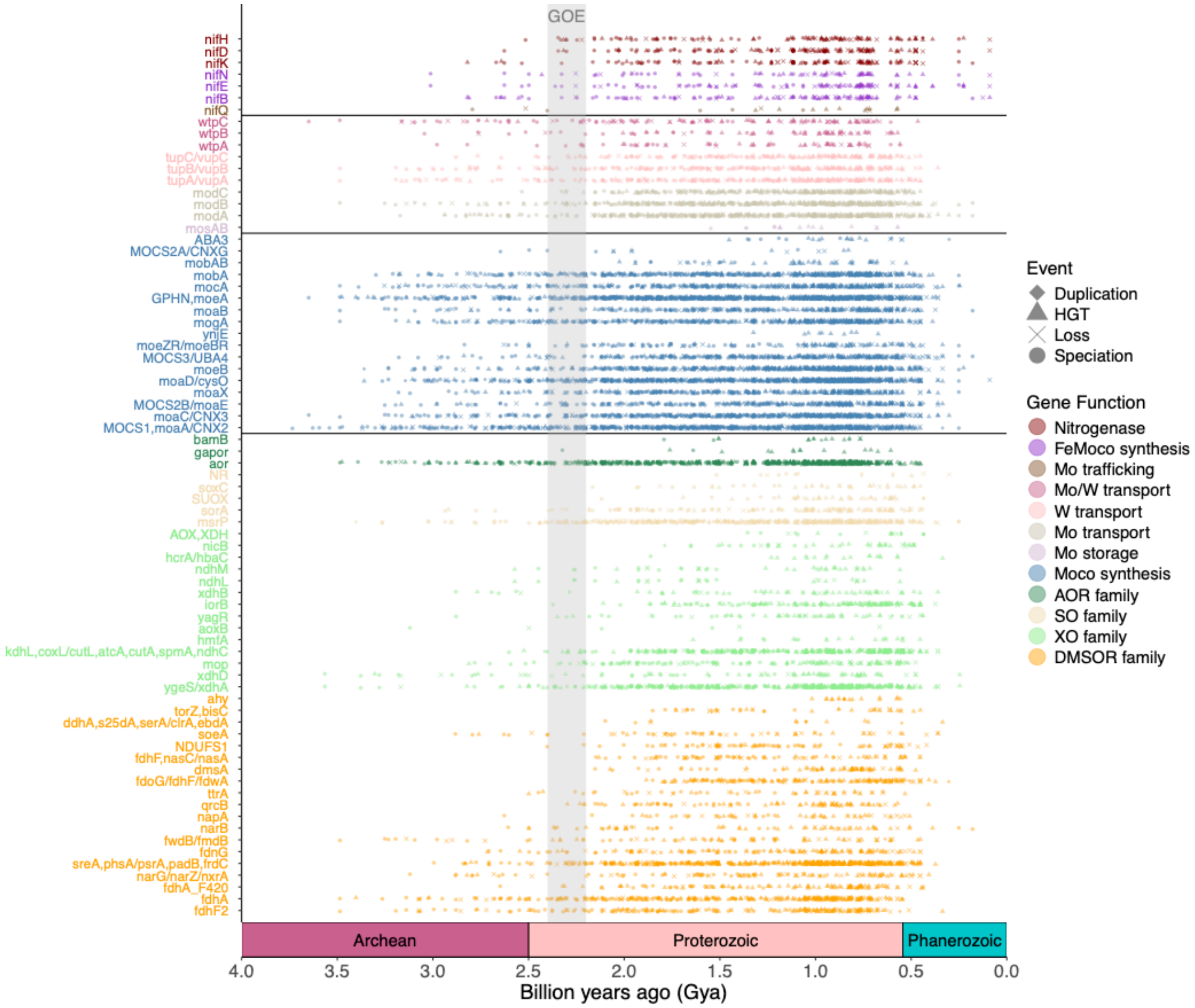


**Figure S11. Gene speciation, duplication, loss, and horizontal gene transfer events from a random reconciliation plotted across time for Mo-/W-related genes as identified by ecceTERA using a new autocorrelated lognormal (LN) clock model from Boden et al. (2024) generated using an earlier estimation for methanogenesis (> 3.46 Ga).** Random reconciliation results (as opposed to symmetric or asymmetric median reconciliations) using a different LN clock model assuming an earlier estimate for methanogenesis from Boden et al. (2024) and the ecceTERA default horizontal gene transfer (HGT) cost of 3 are shown above. Each point represents a single gene event where shapes distinguish different gene event types. Only the midpoint date is shown, except for speciation events where the right node date is shown. Events are colored based on gene groups (shown in boxes above). Lines separate the major Mo-enzyme families, enzymes involved in Moco biosynthesis, metal transport/storage proteins, and proteins involved in nitrogenase assembly and function. The gray rectangle marks an estimated time range for the Great Oxidation Event (GOE, ~2.4-2.2 Gya). Note that gene losses (X's) suggest a gene is no longer being used by a lineage and should not be interpreted as evidence of gene usage.

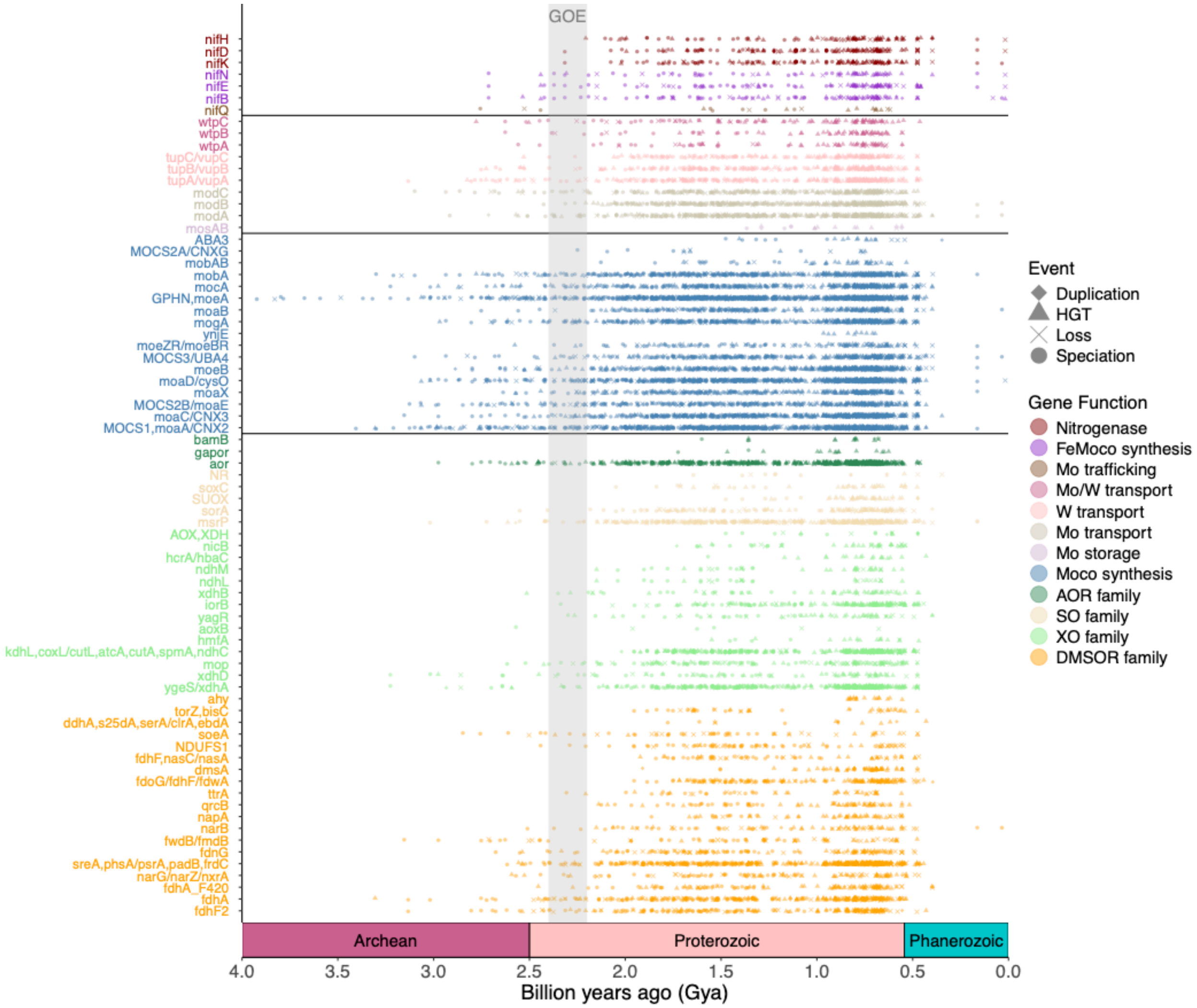


**Figure S12. Gene speciation, duplication, loss, and horizontal gene transfer events from a random reconciliation plotted across time for Mo-/W-related genes as identified by ecceTERA using the uncorrelated gamma multipliers (UGAM) clock model.** Random reconciliation results (as opposed to symmetric or asymmetric median reconciliations) using the UGAM clock model and the ecceTERA default horizontal gene transfer (HGT) cost of 3 are shown above. Each point represents a single gene event where shapes distinguish different gene event types. Only the midpoint date is shown, except for speciation events where the right node date is shown. Events are colored based on gene groups (shown in boxes above). Lines separate the major Mo-enzyme families, enzymes involved in Moco biosynthesis, metal transport/storage proteins, and proteins involved in nitrogenase assembly and function. The gray rectangle marks an estimated time range for the Great Oxidation Event (GOE, ~2.4-2.2 Gya). Note that gene losses (X's) suggest a gene is no longer being used by a lineage and should not be interpreted as evidence of gene usage.

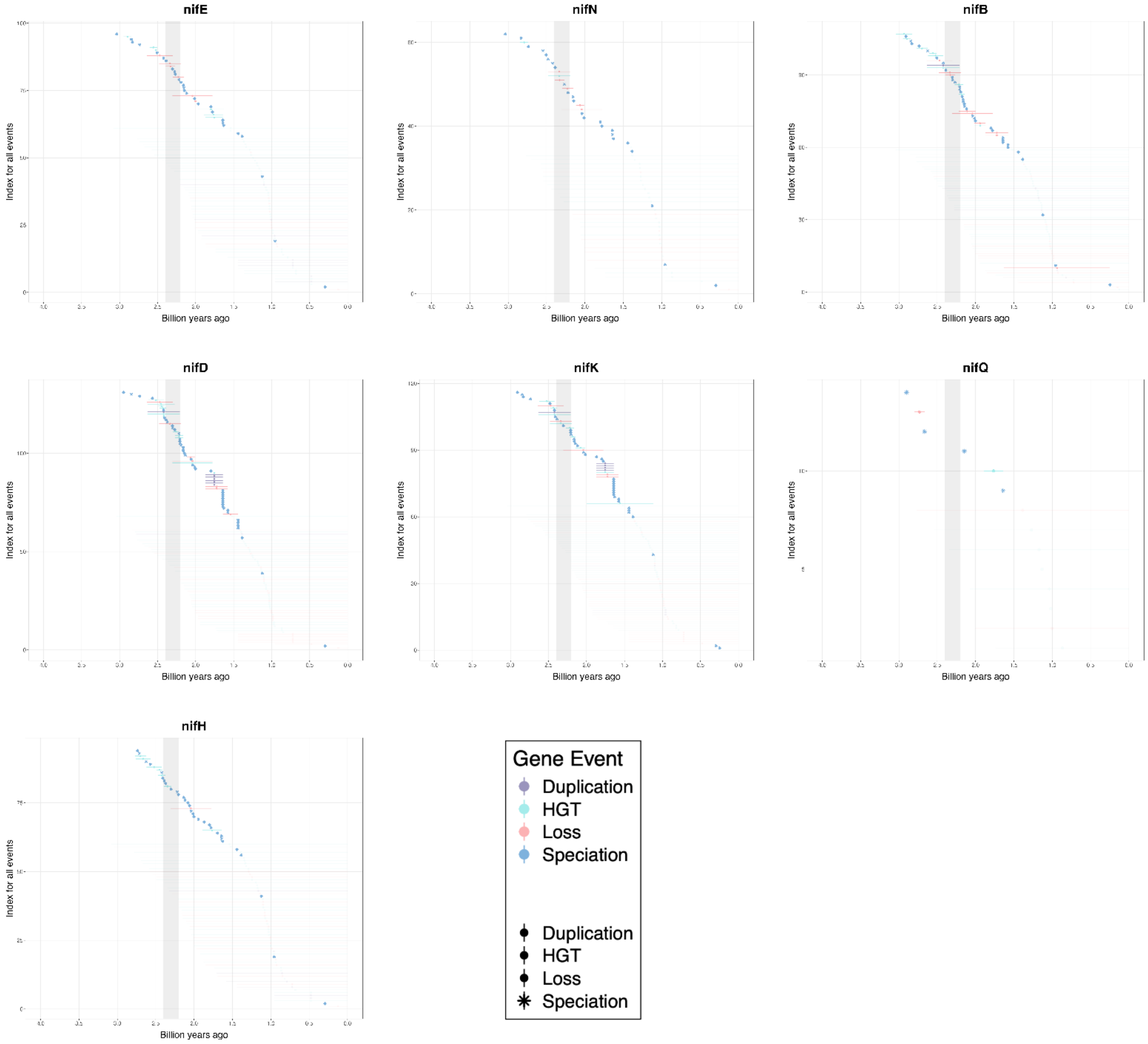


**Figure S13. Time ranges for each gene event (duplication, transfer, loss, and speciation) identified for nitrogenase (*nifHDK*), FeMoco synthesis (*nifNEB*), and nitrogenase Mo trafficking genes (*nifQ*).** Time ranges correspond to branches on a time-calibrated tree of life, where a gene event is estimated to have occurred anywhere along the branch based on symmetric reconciliation results. Lower and upper bounds for ranges correspond to left node and right node dates on the branch a given event occurred on. Only right node dates are shown for speciation events. Events that occurred on terminal branches are shown in faded color, while events occurring on internal branches are shown in solid color. Reconciliations were performed using the CIR clock model with an HGT cost of 3 (default for ecceTERA). Points and associated ranges are colored by the type of gene event (blue = duplication, cyan = horizontal gene transfer, pink = loss, purple= speciation). Gray bars in each plot correspond to an estimated time range for the Great Oxidation Event (GOE) at 2.4-2.2 Gya. Plots are shown from left to right in order of age of the oldest gene event's midpoint date for a given gene (going oldest to youngest).

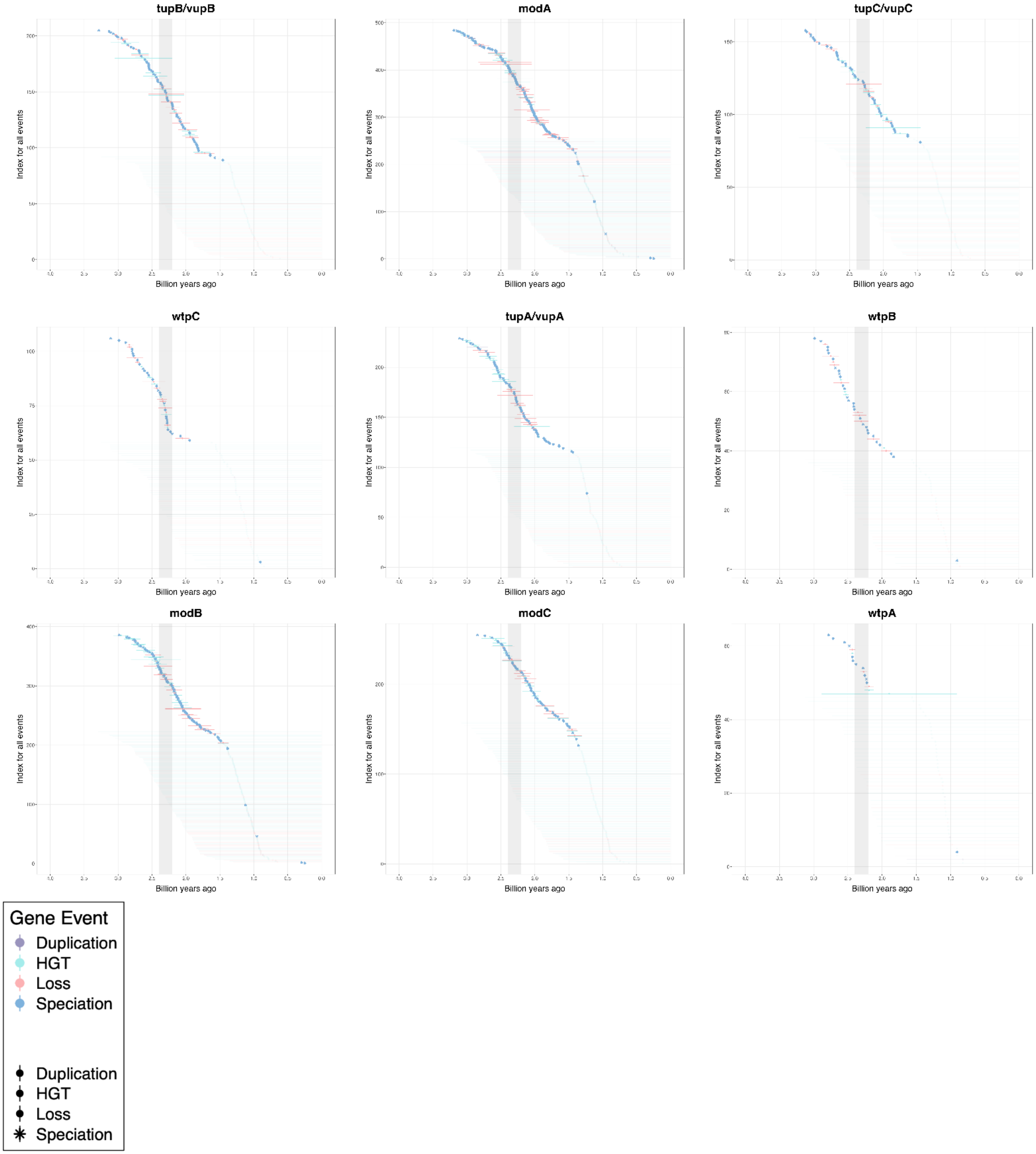

tupB/vupB
modA
tupC/vupC
wtpC
tupA/vupA
wtpB
modB
modC
wtpA
Index for all events
Billion years ago
Gene Event
Duplication
HGT
Loss
Speciation
Duplication
HGT
Loss
Speciation

**Figure S14. Time ranges for each gene event (duplication, transfer, loss, and speciation) identified for high affinity Mo (*modABC*), high affinity W (*tupABC*), and dual-affinity Mo/W (*wtpABC*) transporters.** Time ranges correspond to branches on a time-calibrated tree of life, where a gene event is estimated to have occurred anywhere along the branch based on symmetric reconciliation results. Lower and upper bounds for ranges correspond to left node and right node dates on the branch a given event occurred on. Only right node dates are shown for speciation events. Events that occurred on terminal branches are shown in faded color, while events occurring on internal branches are shown in solid color. Reconciliations were performed using the CIR clock model with an HGT cost of 3 (default for ecceTERA). Points and associated ranges are colored by the type of gene event (blue = duplication, cyan = horizontal gene transfer, pink = loss, purple= speciation). Gray bars in each plot correspond to an estimated time range for the Great Oxidation Event (GOE) at 2.4-2.2 Gya. Plots are shown from left to right in order of age of the oldest gene event's midpoint date for a given gene (going oldest to youngest).

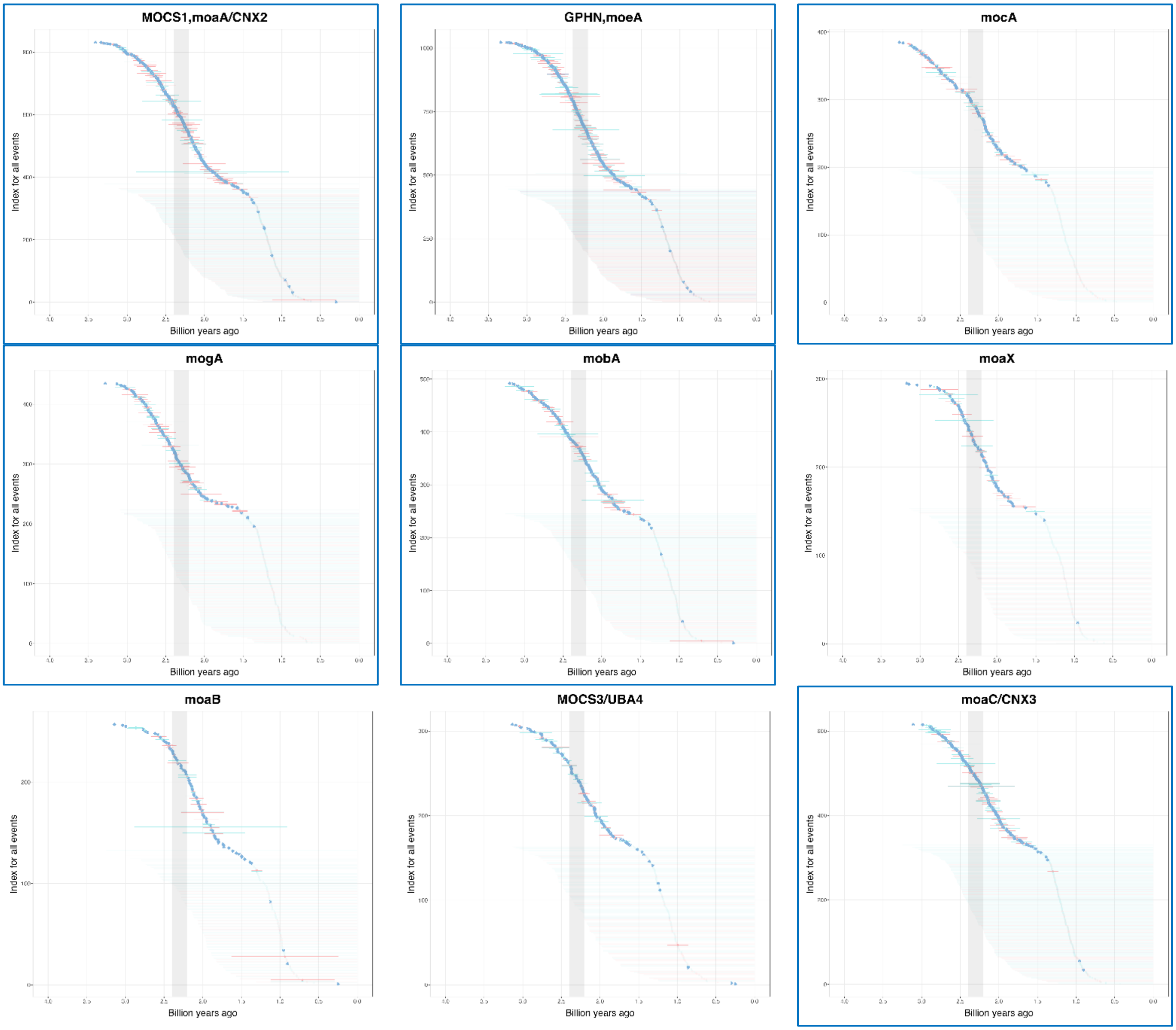
MOCS1,moaA/CNX2
GPHN,moeA
mocA
mogA
mobA
moaX
moaB
MOCS3/UBA4
moaC/CNX3
Index for all events
Billion years ago

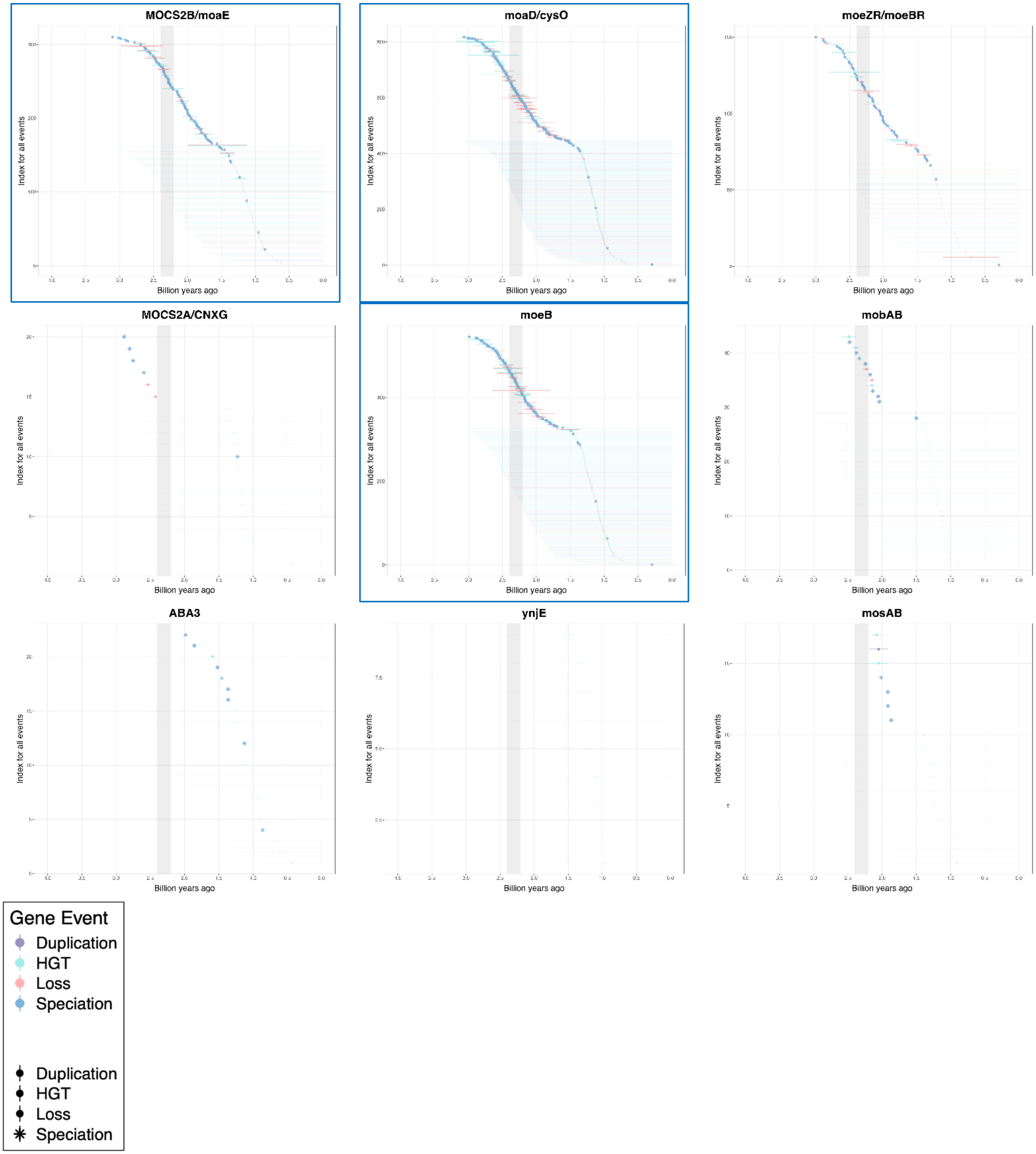


**Figure S15. Time ranges for each gene event (duplication, transfer, loss, and speciation) identified for Moco synthesis genes and Mo storage protein MosAB.** Time ranges correspond to branches on a time-calibrated tree of life, where a gene event is estimated to have occurred anywhere along the branch based on symmetric reconciliation results for most genes and random reconciliation results for a subset (detailed in the methods section) where symmetric results were not available. Lower and upper bounds for ranges correspond to left node and right node dates on the branch a given event occurred on. Only right node dates are shown for speciation events. Events that occurred on terminal branches are shown in faded color, while events occurring on internal branches are shown in solid color. Reconciliations were performed using the CIR clock model with an HGT cost of 3 (default for ecceTERA). Points and associated ranges are colored by the type of gene event (blue = duplication, cyan = horizontal gene transfer, pink = loss, purple= speciation). Gray bars in each plot correspond to an estimated time range for the Great Oxidation Event (GOE) at 2.4-2.2 Gya. Plots are shown from left to right in order of age of the oldest gene event's midpoint date for a given gene (going oldest to youngest). Plots with blue borders correspond to the 9 genes essential for Moco synthesis as described by Mendel & Leimkühler (2015).

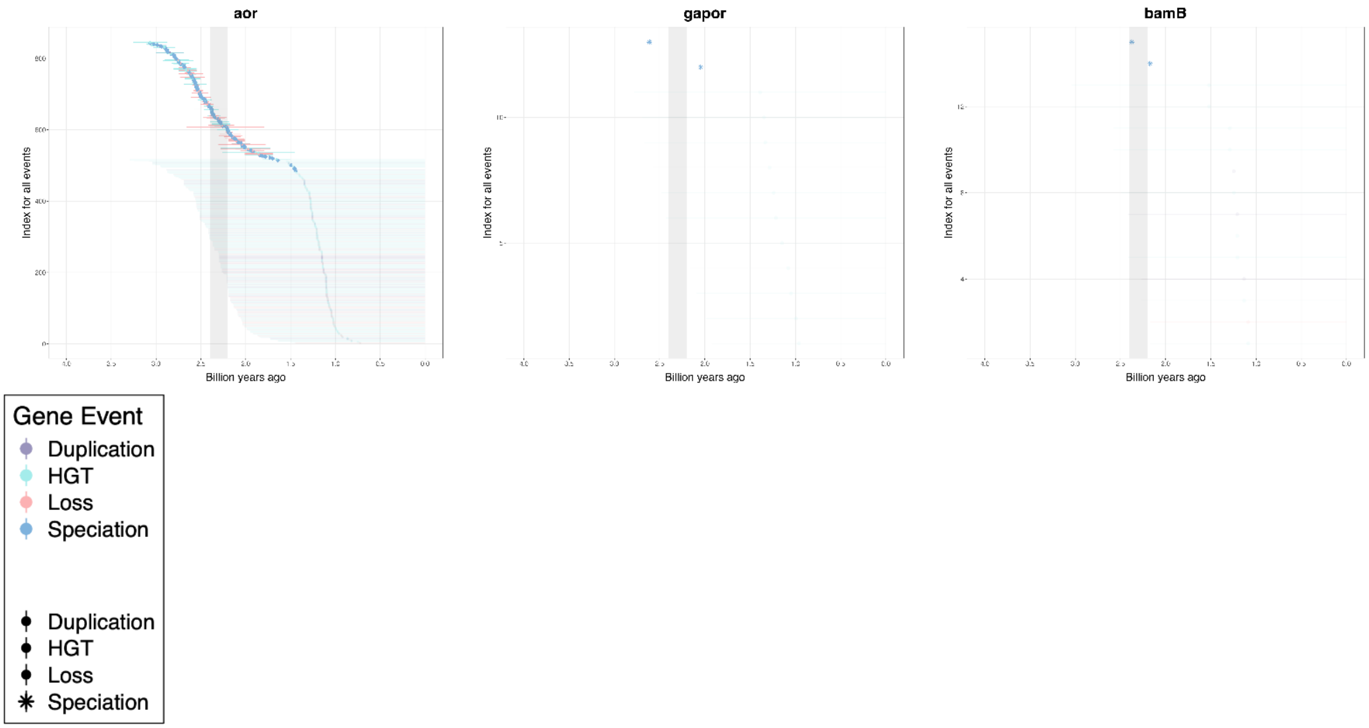


**Figure S16. Time ranges for each gene event (duplication, transfer, loss, and speciation) identified for genes of the aldehyde:ferredoxin oxidoreductase (AOR) family of tungstoenzymes.** Time ranges correspond to branches on a time-calibrated tree of life, where a gene event is estimated to have occurred anywhere along the branch based on symmetric reconciliation results for most genes and random reconciliation results for a subset (detailed in the methods section) where symmetric results were not available. Lower and upper bounds for ranges correspond to left node and right node dates on the branch a given event occurred on. Reconciliations were performed using the CIR clock model with an HGT cost of 3 (default for ecceTERA). Points and associated ranges are colored by the type of gene event (blue = duplication, cyan = horizontal gene transfer, pink = loss, purple= speciation). Gray bars in each plot correspond to an estimated time range for the Great Oxidation Event (GOE) at 2.4-2.2 Gya. Plots are shown from left to right in order of age of the oldest gene event's midpoint date for a given gene (going oldest to youngest).

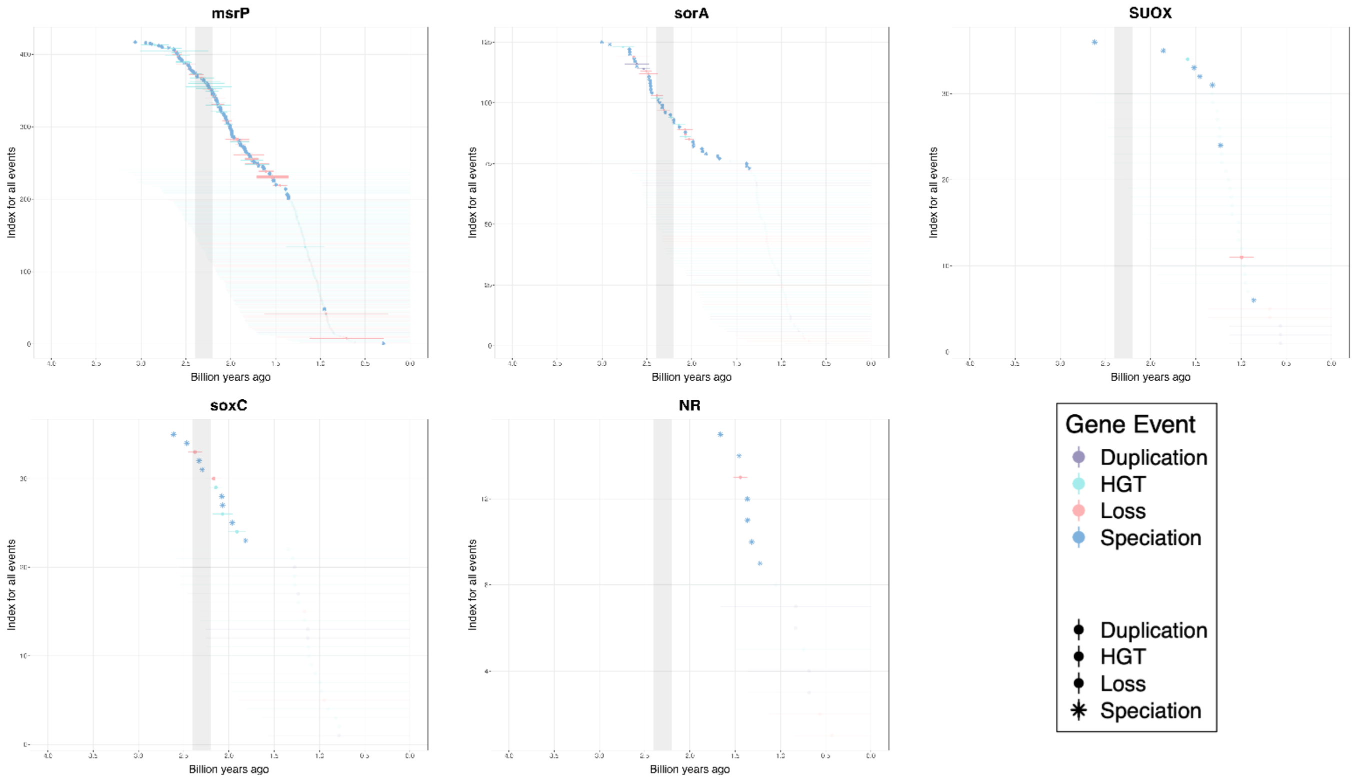


**Figure S17. Time ranges for each gene event (duplication, transfer, loss, and speciation) identified for the sulfite oxidase (SO) family of molybdoenzymes.** Time ranges correspond to branches on a time-calibrated tree of life, where a gene event is estimated to have occurred anywhere along the branch based on symmetric reconciliation results. Lower and upper bounds for ranges correspond to left node and right node dates on the branch a given event occurred on. Reconciliations were performed using the CIR clock model with an HGT cost of 3 (default for ecceTERA). Points and associated ranges are colored by the type of gene event (blue = duplication, cyan = horizontal gene transfer, pink = loss, purple= speciation). Gray bars in each plot correspond to an estimated time range for the Great Oxidation Event (GOE) at 2.4-2.2 Gya. Plots are shown from left to right in order of age of the oldest gene event's midpoint date for a given gene (going oldest to youngest).

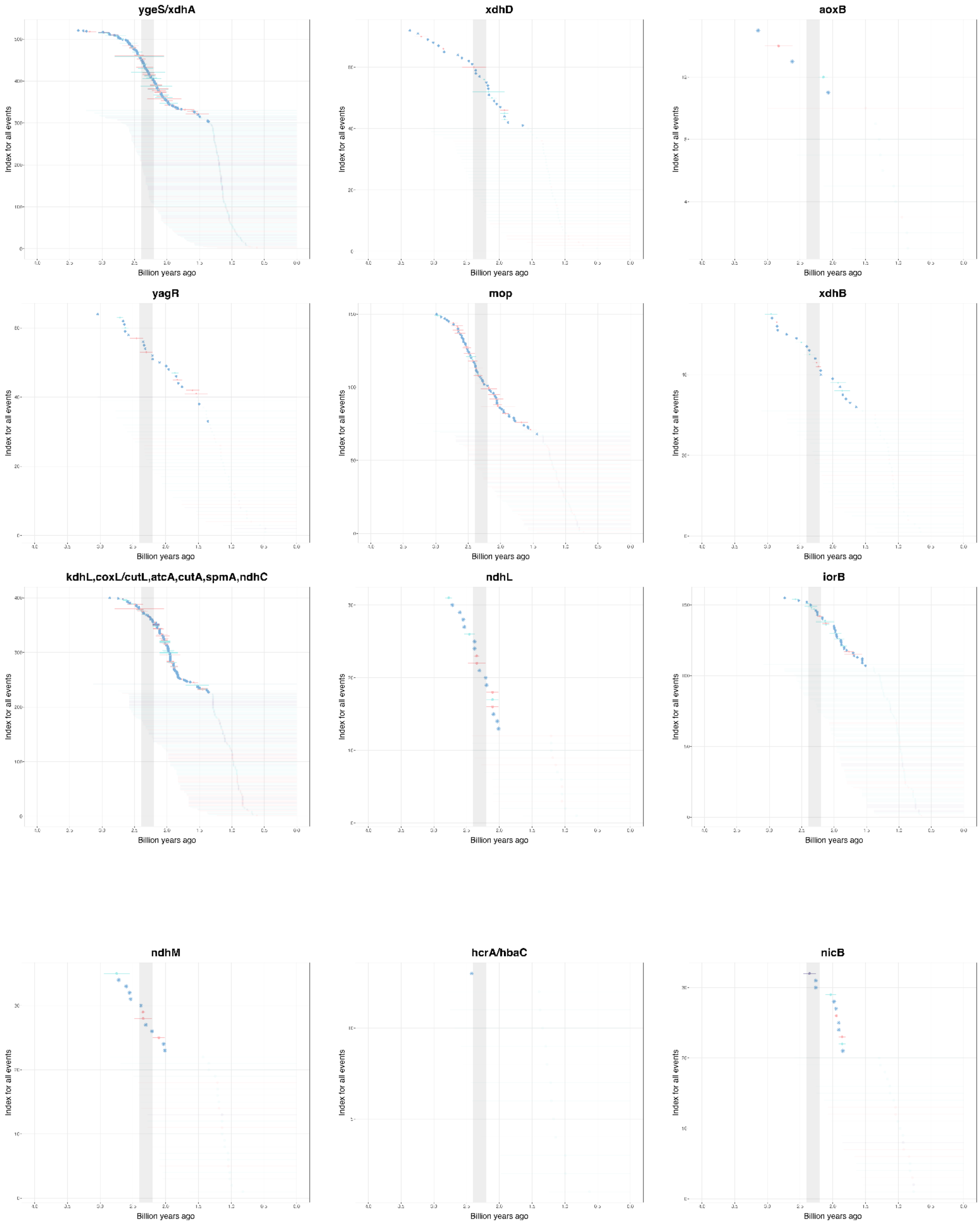
ygeS/xdhA
xdhD
aoxB
yagR
mop
xdhB
kdhL,coxL/cutL,atcA,cutA,spmA,ndhC
ndhL
iorB
ndhM
hcrA/hbaC
nicB
Index for all events
Billion years ago

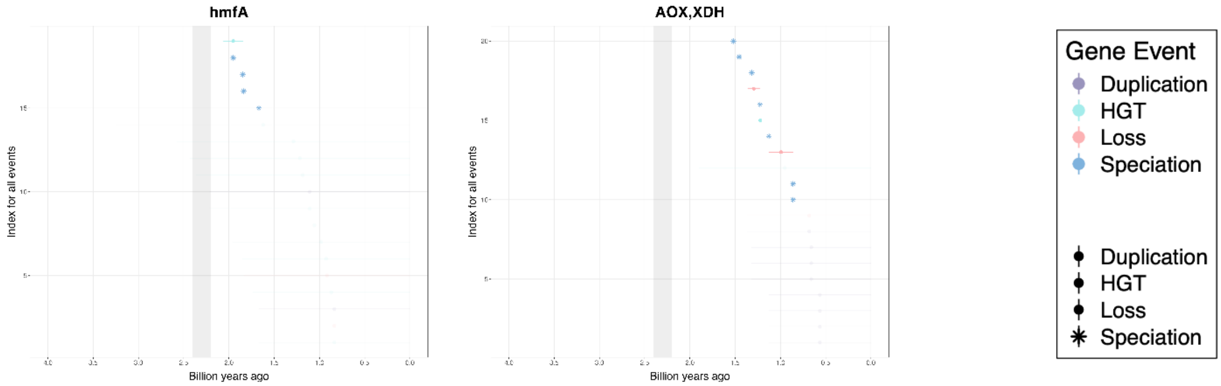


**Figure S18. Time ranges for each gene event (duplication, transfer, loss, and speciation) identified for the xanthine oxidase (XO) family of molybdoenzymes.** Time ranges correspond to branches on a time-calibrated tree of life, where a gene event is estimated to have occurred anywhere along the branch based on symmetric reconciliation results for most genes and random reconciliation results for a subset (detailed in the methods section) where symmetric results were not available. Lower and upper bounds for ranges correspond to left node and right node dates on the branch a given event occurred on. Reconciliations were performed using the CIR clock model with an HGT cost of 3 (default for ecceTERA). Points and associated ranges are colored by the type of gene event (blue = duplication, cyan = horizontal gene transfer, pink = loss, purple= speciation). Gray bars in each plot correspond to an estimated time range for the Great Oxidation Event (GOE) at 2.4-2.2 Gya. Plots are shown from left to right in order of age of the oldest gene event's midpoint date for a given gene (going oldest to youngest).

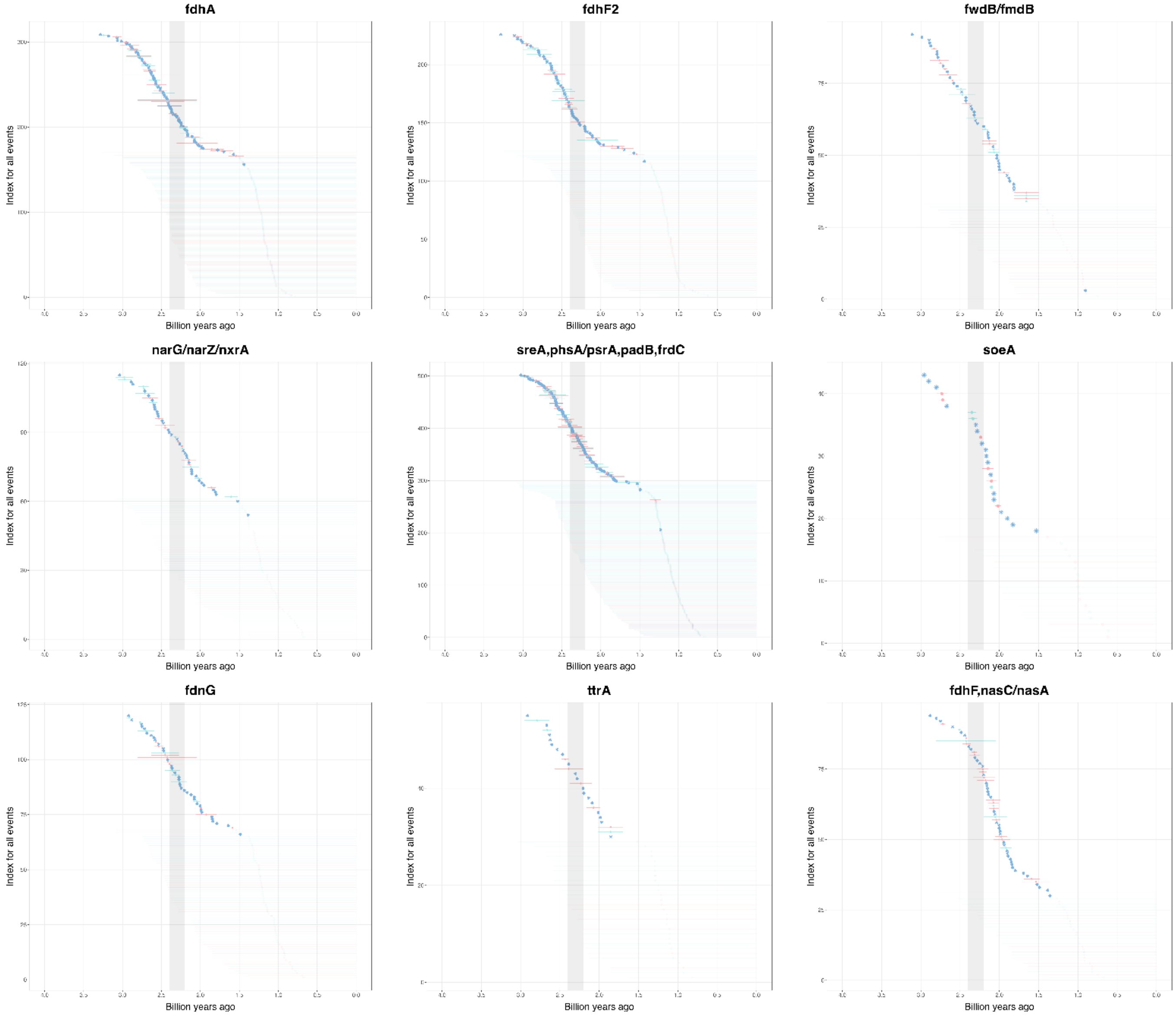

fdhA
fdhF2
fwdB/fmdB
narG/narZ/nxrA
sreA,phsA/psrA,padB,frdC
soeA
fdnG
ttrA
fdhF,nasC/nasA
Index for all events
Billion years ago

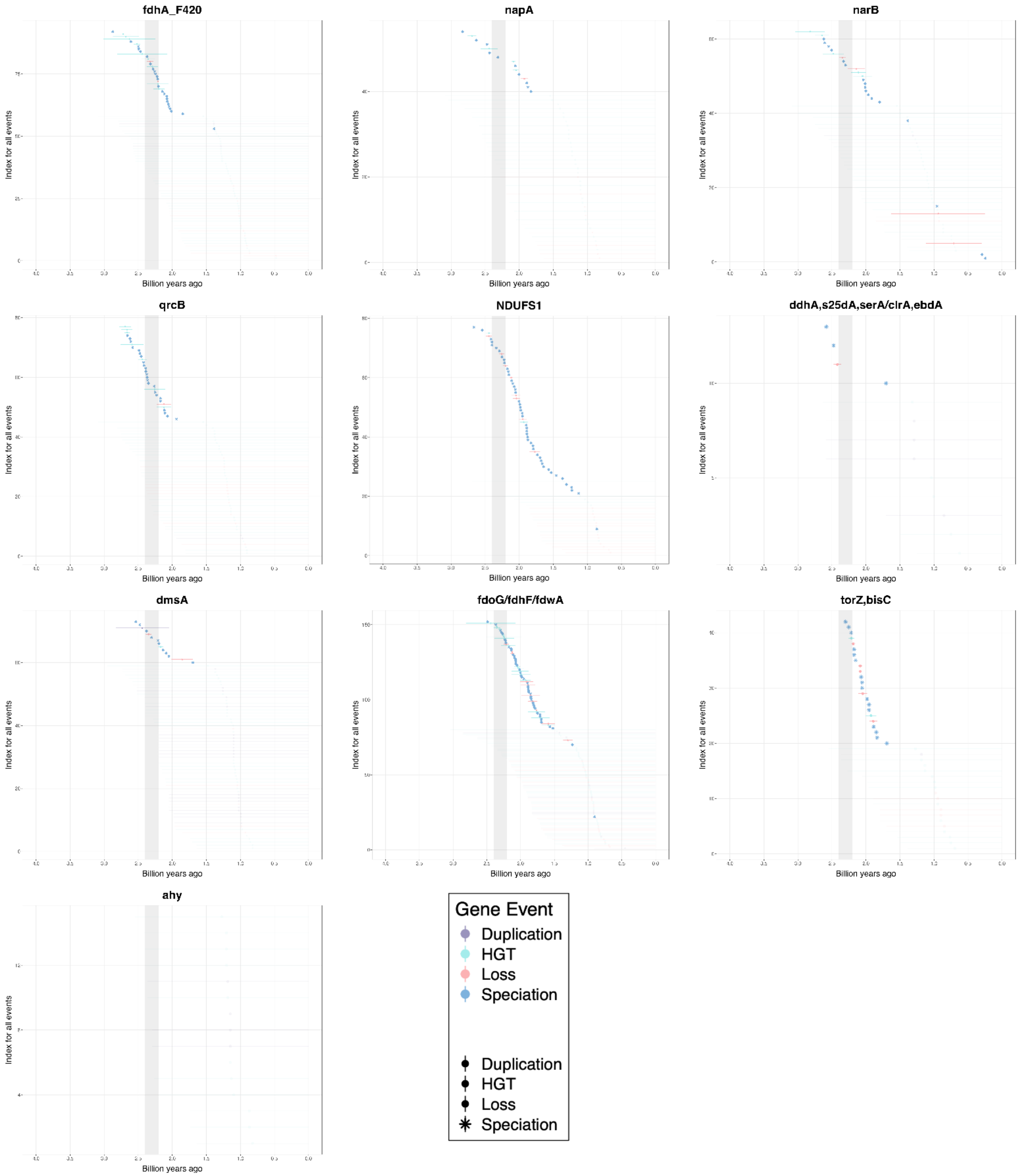


**Figure S19. Time ranges for each gene event (duplication, transfer, loss, and speciation) identified for the dimethylsulfide reductase (DMSOR) family of molybdoenzymes.** Time ranges correspond to branches on a time-calibrated tree of life, where a gene event is estimated to have occurred anywhere along the branch based on symmetric reconciliation results for most genes and random reconciliation results for a subset (detailed in the methods section) where symmetric results were not available. Lower and upper bounds for ranges correspond to left node and right node dates on the branch a given event occurred on. Reconciliations were performed using the CIR clock model with an HGT cost of 3 (default for ecceTERA). Points and associated ranges are colored by the type of gene event (blue = duplication, cyan = horizontal gene transfer, pink = loss, purple= speciation). Gray bars in each plot correspond to an estimated time range for the Great Oxidation Event (GOE) at 2.4-2.2 Gya. Plots are shown from left to right in order of age of the oldest gene event's midpoint date for a given gene (going oldest to youngest).

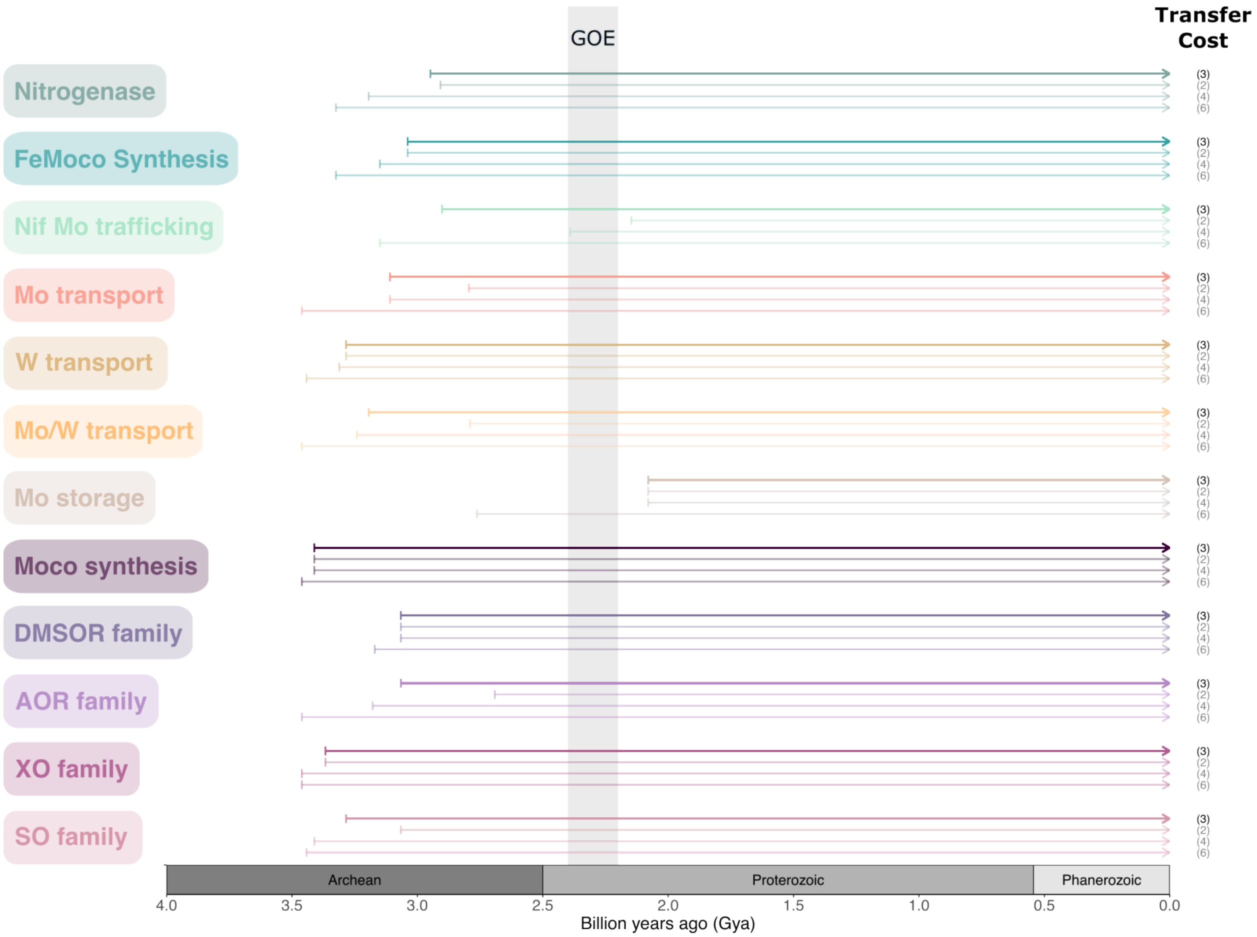


**Figure S20. Arrows depicting earliest gene events for a given group of Mo-/W-related enzymes based on reconciliations using different horizontal gene transfer (HGT) costs.** Starting points of arrows represent the midpoint date of the earliest gene event among genes within a given group, based on reconciliations using the CIR clock model with HGT costs of 3 (ecceTERA default), 2, 4, and 6 in order from top to bottom in the plot. Arrows are colored by gene group. The gray bar from 2.2 to 2.4 Gya corresponds to the estimated time range for the Great Oxidation Event (GOE).
